# A CDK1 phospho-switch reprograms TRAIP to unload replisomes in mitosis

**DOI:** 10.1101/2024.11.30.626186

**Authors:** Geylani Can, Maksym Shyian, Archana Krishnamoorthy, Samreen Ahmed, Yang Lim, R. Alex Wu, Raphael Pavani, Manal S. Zaher, André Nussenzweig, Markus Räschle, Thomas E. Wilson, Thomas W. Glover, Johannes C. Walter, David Pellman

## Abstract

Cells entering mitosis with incompletely replicated DNA face catastrophic chromosome segregation failure. During interphase, the replisome-associated E3 ubiquitin ligase TRAIP ubiquitylates barriers in front of the fork to allow replisome progression. In mitosis, TRAIP is reprogrammed from a *trans*-acting to a *cis*-acting ligase that can ubiquitylate the replisome itself. This enables the processing of unreplicated DNA by promoting replisome disassembly, fork breakage, and joining of the broken chromosome arms. Here, we describe a mechanism for this reprogramming: the ATPase TTF2 is recruited to the replisome, where its non-catalytic N-terminal domain tethers Cyclin B-CDK1-phosphorylated TRAIP to the leading strand DNA polymerase ε in a geometry that allows replisome ubiquitylation. Thus, a phospho-regulated architectural switch alters replisome organization in mitosis to safeguard genome integrity before chromosome segregation.

## Main Text

Replisomes encounter many obstacles that can disrupt DNA replication and genome integrity (*1*). These include transcription complexes, as well as covalent DNA-protein cross-links (DPCs) and DNA inter-strand cross-links (ICLs) that are generated by endogenous aldehydes and chemotherapeutics. Cells employ various strategies to overcome these challenges. If these strategies fail, cells enter mitosis with under-replicated DNA. This leads to gaps and breaks at difficult-to-replicate loci called common fragile sites (termed CFS expression) and it triggers additional defense mechanisms to prevent chromosomal instability. These defenses include unwinding (dissolution) or breakage (resolution) of the unreplicated segment (*2*). The latter pathway can result in large deletions at CFSs and elsewhere in the genome and involves nucleolytic cleavage of the stalled forks and joining of the resulting broken chromosomes by alternative end-joining (*3*). Although CFS processing can result in a deletion and sister chromatid exchange, this outcome avoids the potentially catastrophic consequences of chromosome non-disjunction.

The E3 ubiquitin ligase TRAIP has been linked to DNA repair in interphase and to the response to under-replicated DNA in mitosis (*4*). TRAIP is essential for embryonic development and cell proliferation (*5*)(*6*)(*7*), and hypomorphic mutations in TRAIP are associated with dwarfism in humans (*8*). Extensive evidence suggests that TRAIP plays a central role in overcoming replication obstacles in S phase (*4*). Cells lacking TRAIP are sensitive to agents that induce DNA inter-strand crosslinks (ICLs) and DNA-protein crosslinks (DPCs), and to conditions that promote replication-transcription collisions (*9*). Indirect evidence suggests that TRAIP binds replisomes with its catalytic RING domain directed ahead of the CMG, allowing it to ubiquitylate “*in trans*” any protein barrier encountered by the replisome (e.g. DPCs), while being unable to ubiquitylate “*in cis*” the replisome with which it travels (fig. S1A; “hood ornament” model)(*10*, *11*). This block to *cis* ubiquitylation is crucial to avoid unscheduled CMG unloading during S phase, which would lead to catastrophic fork arrest and genome instability (*12*)(*13*).

TRAIP’s function at the replisome appears to change fundamentally in mitosis (fig. S1B). Thus, when replisomes stall on either side of a lac repressor (LacR) array in mitotic egg extracts, TRAIP acquires the ability to ubiquitylate the stalled CMG with which it travels, leading to p97-dependent CMG unloading, fork breakage, and DNA polymerase θ (pol θ)-mediated alternative end-joining (TMEJ; fig. S1C) (*14*, *15*). Consistent with these observations, when CMG unloading in S phase is prevented in cells by neutralizing CRL2^Lrr1^, the E3 ligase that normally promotes replication termination, CMGs are unloaded in mitosis by TRAIP (*14*)(*16*). Furthermore, when the completion of DNA replication is experimentally impaired, TRAIP promotes mitotic DNA synthesis, and it suppresses the formation of anaphase bridges and micronuclei (*17*). Together with other results (*18*), these observations suggest that when forks fail to fully converge in S or G2, TRAIP-mediated symmetrical fork cleavage in mitosis could restore one intact chromosome and regenerate the other via alternative end joining, leading to a deletion and sister chromatid exchange (fig. S1C) (*14*). This model accounts for key aspects of common fragile site expression and the sequence features associated with common fragile site rearrangements in cancer (*3*). The TRAIP-dependent unloading of replisomes in mitosis appears to be a last-ditch cellular response to avert anaphase with unreplicated DNA. How TRAIP changes its specificity between interphase and mitosis is unclear.

The SWI/SNF ATPase TTF2 was previously shown to promote the eviction of RNAPII from mitotic chromosomes (*19*). TTF2 is a cytoplasmic protein that only gains access to chromosomes when the nuclear envelope breaks down at the beginning of mitosis. Here, we report an unexpected role for TTF2 in activating the mitotic function of TRAIP. Using *Xenopus* egg extracts, we show that the N-terminal Zinc fingers of TTF2 interact with TRAIP that has been phosphorylated by B-CDK1. In addition, a short TTF2 peptide that resides adjacent to the TTF2 zinc fingers binds to the POLE2 subunit of DNA polymerase ε. These two TTF2 contacts are sufficient to tether TRAIP to the replisome in a geometry that allows mitotic CMG ubiquitylation and unloading, fork breakage, and alternative end-joining. In mammalian cells, disruption of the TRAIP-TTF2 complex impairs mitotic CMG unloading and modestly reduces deletions at common fragile sites. On the other hand, TTF2’s ATPase domain is sufficient to promote RNAPII eviction, indicating that TTF2 uses distinct mechanisms to remove replication and transcription complexes from mitotic chromosomes. Together, our results identify a structural and functional remodeling of TRAIP on the replisome that is driven by TTF2 and CDK1.

## Results

### Identification of CDK sites that promote mitotic TRAIP function

We hypothesized that mitotic TRAIP activation involves its phosphorylation by Cyclin B-CDK1 (B-CDK1). To test this idea, we used frog egg extracts because they faithfully recapitulate TRAIP’s interphase role in ubiquitylating replication barriers and its mitotic role in replisome ubiquitylation and disassembly (*10*, *14*, *15*, *20*). TRAIP forms a dimer via its coiled-coil and leucine zipper domains (predictomes.org; (*21*)), and it contains three conserved B-CDK1 consensus S/TP sites (Fig. 1A; red residues; fig. S2A).

**Fig. 1.**
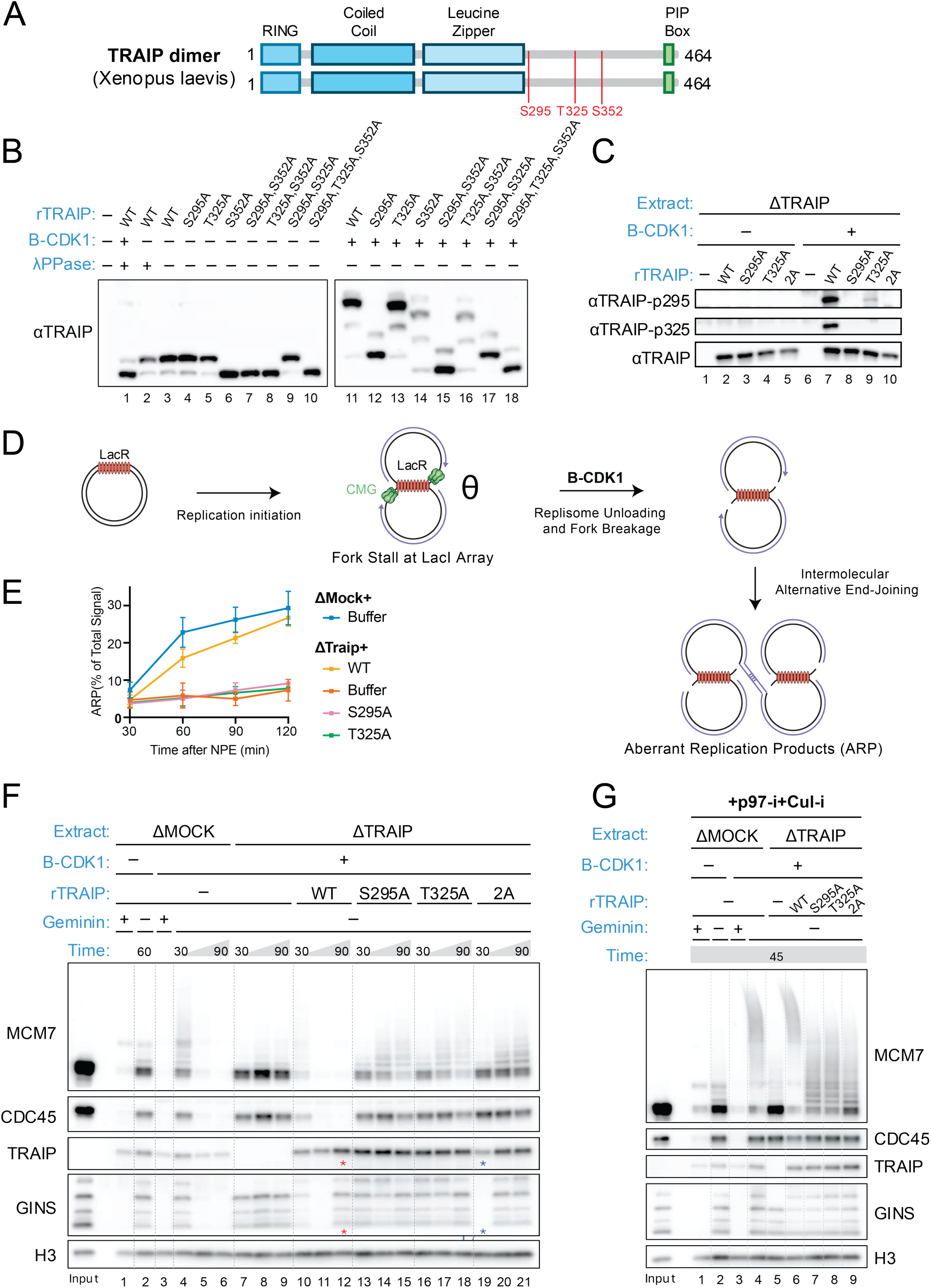
TRAIP is activated by B-CDK1 phosphorylation in mitotic egg extracts. **(A)** Domain organization of *Xenopus laevis* TRAIP, which is a dimer (*21*). The RING, coiled-coil, leucine zipper, and PIP box are indicated. Conserved CDK consensus sites S295, T325, and S352 are shown in red. **(B)** TRAIP is phosphorylated by B-CDK1. The indicated recombinant *Xenopus* TRAIP proteins were expressed in wheat germ transcription-translation extracts (TTE) and added to TRAIP-depleted *Xenopus* nucleoplasmic extract (NPE) optionally supplemented with B-CDK1. Where indicated, samples were treated with λ phosphatase, separated on a Phos-tag gel, and blotted for TRAIP protein. **(C)** S295 and T325 are major B-CDK1 phosphorylation sites. The indicated *Xenopus* TRAIP proteins were added to TRAIP-depleted NPE that was optionally supplemented with B-CDK1. Samples were blotted with the indicated antibodies, including two phospho-specific TRAIP antibodies, whose specificity was validated by the absence of signal in the appropriate phospho-site mutations (lanes 8 and 9). 2A, TRAIP S295A/T325A double mutant. **(D)** Model for TRAIP-dependent processing of stalled replication forks in mitosis. After fork stalling at a LacR array, Cyclin B-CDK1 triggers TRAIP-dependent CMG ubiquitylation and replisome unloading, leading to fork breakage and formation of aberrant replication products (ARP) by alternative end-joining. **(E)** TRAIP’s CDK sites are required for ARP formation. A plasmid with 48 tandem *lacO* sites bound to LacR (“LacR plasmid”) was replicated in the indicated egg extracts supplemented with [α-^32^P]dATP and different TRAIP mutants expressed in TTE. Samples were withdrawn at 30, 45, 60, 90, and 180 minutes after replication was initiated and subjected to gel electrophoresis and autoradiography. ARP formation was quantified over time after addition of NPE and plotted as percentage of total replication products. A representative gel is shown in fig. S2B. **(F)** TRAIP’s CDK sites are required for CMG unloading from stalled forks. LacR plasmid was replicated in the indicated egg extracts supplemented with different TRAIP mutants expressed in TTE, geminin (to prevent licensing and replication initiation), and B-CDK1, as noted. At 30, 45, and 90 minutes, chromatin was recovered and analyzed by blotting with the indicated antibodies. Two chromatin samples that were run on the same gel and blotted for TRAIP and GINS were inadvertently switched (red and blue asterisks). Here and in other complicated gels, samples with related conditions are grouped by dotted lines. **(G)** TRAIP’s CDK sites are required for ubiquitylation of stalled CMGs. LacR plasmid was replicated and recovered as in (F), except extracts contained NMS873 (p97-i) and MLN4924 (Cul-i; to inhibit CRL2^Lrr1^-dependent ubiquitylation), and chromatin was recovered at 45 minutes.

To address whether TRAIP is phosphorylated in egg extracts, we added TRAIP that was expressed in a transcription-translation extract (TTE) to TRAIP-depleted frog egg extracts that were also supplemented with B-CDK1 and separated the proteins on a Phos-tag gel (*22*). As shown in Fig. 1B, TRAIP underwent a large mobility shift in the presence of B-CDK1 (compare lanes 3 and 11). Mutating serine 352 to alanine increased the mobility of all forms of TRAIP in interphase and mitotic egg extract (Fig. 1B, lanes 6 and 14), indicating that this site is constitutively phosphorylated throughout the cell division cycle. Accordingly, S352 corresponds to an ideal CDK phosphorylation site (SPTK) that could be phosphorylated by CDK2 in both extracts. In contrast, mutating S295 or T325 reduced TRAIP mobility in mitosis but not in interphase, with S295A having a dramatic effect (Fig. 1B, lanes 3 and 4 vs. 11 and 12) and T325A having a small, but readily detectable effect (Fig. 1B, lanes 3 and 5 vs. 11 and 13). To verify that S295 and T325 were phosphorylated, we generated phospho-specific antibodies (Fig. 1C, lanes 7-9) and confirmed that both residues were indeed phosphorylated only in extracts supplemented with B-CDK1 (Fig. 1C, lanes 2 vs. 7), but not with Cyclin A2-CDK2 or Cyclin A2-CDK1, even when A-CDKs were used at higher concentrations than B-CDK1 (fig. S2E). Interestingly, mutating S295 abolished phosphorylation at both sites (Fig. 1C, lane 8), whereas mutating T325 greatly reduced but did not eliminate S295 phosphorylation (Fig. 1C, lane 9). These results suggest that B-CDK1 phosphorylates both sites, and that S295 phosphorylation primes T325 phosphorylation whereas S295 can be phosphorylated in the absence of T325 phosphorylation, albeit inefficiently.

We next addressed the potential function of these phosphorylation events. We showed previously that when a plasmid containing a lac repressor array (LacR plasmid) is replicated in frog egg extract containing B-CDK1, replisomes stall at the outer edges of the array, generating a “θ” structure (Fig. 1D)(*14*). TRAIP then promotes CMG ubiquitylation, which leads to replisome disassembly by the p97 ATPase, followed by fork breakage. The broken forks undergo pol θ-mediated end-joining to generate complex, <u>a</u>berrant <u>r</u>eplication <u>p</u>roducts (ARPs) that are retained in the well of the gel (Fig. 1D and fig. S2B and C). As expected, TRAIP depletion greatly reduced, and re-addition of TRAIP^WT^ largely rescued ARP formation (Fig. 1E). In contrast, neither TRAIP^S295A^ nor TRAIP^T325A^ supported ARP formation (Fig. 1E; fig. S2B and C), establishing that these CDK sites are required for TRAIP’s mitotic role. Consistent with defective ARP formation, the unloading of stalled CMGs that is normally promoted by TRAIP^WT^ in mitotic extracts (Fig. 1F, lanes 10-12)(*14*) was not supported by TRAIP^S295A^ or TRAIP^T325A^ (Fig. 1F, lanes 13-18; fig. S2C). Accordingly, MCM7 ubiquitylation (measured in the presence of an inhibitor of p97 to prevent CMG unloading) was also greatly diminished in the presence of the phospho-site mutations (Fig. 1G, lanes 6-8; fig. S2D), probably below the threshold normally required for CMG unloading (*23*)(*10*). Mutating S295 and T325 together did not further reduce CMG ubiquitylation compared to the single mutants (Fig. 1G, lanes 7-9; fig. S2D), indicating that the two phosphorylation events control the same pathway. In contrast, TRAIP depletion reduced CMG ubiquitylation still further (Fig. 1G, lane 5), suggesting that a basal level of TRAIP-dependent CMG ubiquitylation still occurs in the absence of these phosphorylation events, perhaps due to residual TRAIP binding to the replisome. However, this residual activity was not sufficient to support efficient CMG unloading (Fig. 1F, lanes 13-21). Consistent with our results, simultaneous mutation of T280, S295, and S352 abolishes mitotic TRAIP activity (*24*).

Importantly, like TRAIP^WT^ (*10*), TRAIP^S295A^ and TRAIP^T325A^ supported the unloading of CMGs that have converged on an ICL in both interphase and mitotic egg extracts (fig. S3A-C), indicating that these mutations still support “*in trans*” ubiquitylation of converged CMGs. Thus, B-CDK1 phosphorylation of TRAIP on at least two sites is specifically required for mitotic *in cis* ubiquitylation of CMG by TRAIP, whereas CDK phosphorylation does not inhibit TRAIP’s capacity to ubiquitylate barriers *in trans*, such as converged replisomes, in interphase or mitosis.

### TTF2 is essential for TRAIP-dependent CMG unloading

To identify new proteins that cooperate with TRAIP in mitosis and to explain how TRAIP is activated by B-CDK1, we screened for proteins that specifically associate with mitotic stalled replisomes. To this end, we replicated LacR plasmids in egg extract containing B-CDK1 and p97 inhibitor to accumulate mitotic replisomes. We then isolated chromatin and performed mass spectrometry. As controls, we recovered chromatin from reactions lacking p97-i (mitosis with replisome removal) or B-CDK1 (interphase without replisome removal). This analysis identified a handful of proteins that were only enriched on mitotic chromatin containing replisomes (Fig. 2A, red box and fig. S4). TTF2, which contains a C-terminal SWI/SNF ATPase domain (Fig. 2B), was particularly interesting because it removes RNA polymerase II from mitotic chromosomes (*19*), a process reminiscent of mitotic CMG eviction. Interestingly, TTF2 is cytoplasmic during interphase and only gains access to chromosomes after mitotic nuclear envelope breakdown (*19*)(*25*).

**Fig. 2.**
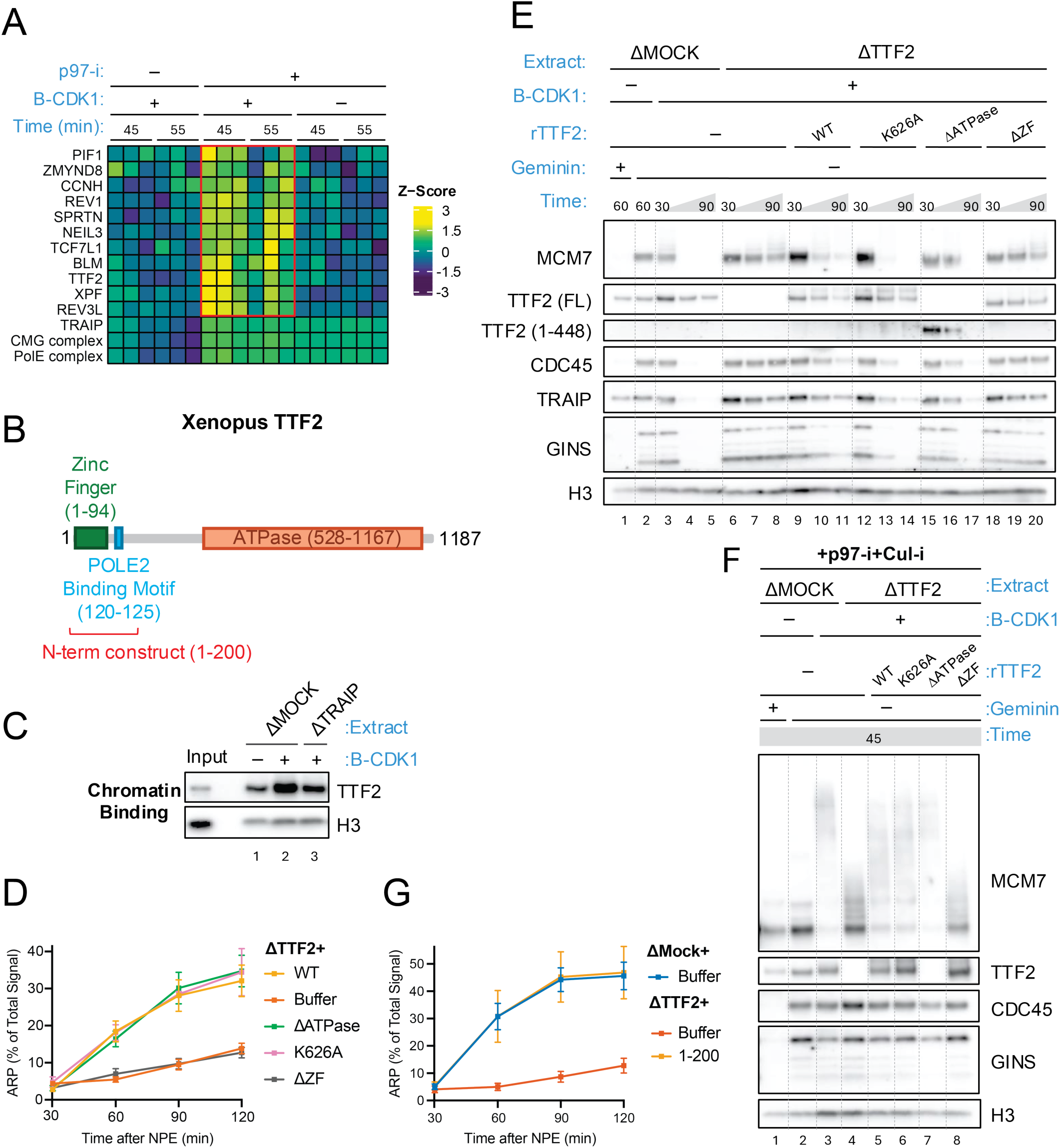
TTF2’s zinc finger domain is required for mitotic CMG ubiquitylation by TRAIP. **(A)** Mass spectrometry analysis of chromatin-associated proteins. LacR plasmid was replicated in the indicated egg extracts (as in Fig. 1E), recovered, and analyzed by label-free mass spectrometry. Three replisome components are shown (bottom three rows), as are proteins specifically enriched in the presence of both B-CDK1 and p97-i (red box). Full datasets and statistical analysis are provided in fig. S4 and Table S1. **(B)** Domain organization and key elements of *Xenopus* TTF2. **(C)** TRAIP is required for mitotic TTF2 recruitment. LacR plasmid replicated in the indicated extracts (as in Fig. 1E) was recovered (as in Fig. 1F) after 30 minutes, and blotted for TTF2 and histone H3. **(D)** The TTF2 Zinc finger domain but not its ATPase function is required for ARP formation. ARP formation was quantified over time after NPE addition in extracts supplemented with the indicated TTF2 constructs and plotted as a percentage of total replication products. **(E)** The TTF2 Zinc finger domain is required for stalled CMG unloading in mitosis. CMG unloading was measured (as in Fig. 1F) using the indicated extracts and TTF2 constructs. **(F)** The TTF2 Zinc finger domain is required for CMG ubiquitylation. CMG ubiquitylation was performed (as in Fig. 1G) using the indicated extracts and TTF2 constructs. **(G)** TTF2^1-200^ is sufficient to support ARP formation. ARP formation was quantified as in Fig. 1E in extracts supplemented with the indicated TTF2 fragments.

We raised a peptide antibody against *Xenopus* TTF2 (fig. S5A) and found that the protein’s recruitment to chromatin was enhanced in mitotic extract, in a manner that partially depended on TRAIP (Fig. 2C). Moreover, like TRAIP depletion, depletion of TTF2 from egg extract greatly reduced ARP formation (Fig. 2D, fig. S5B), CMG unloading (Fig. 2E, lanes 3-8), and CMG ubiquitylation (Fig. 2F, lanes 3-4), and these defects were rescued with recombinant wild type TTF2 (Fig. 2D-F; fig. S5B). Therefore, TTF2 is essential for TRAIP-dependent CMG unloading in mitotic egg extract.

We next determined which domains of TTF2 promote CMG unloading. TTF2 contains a pair of N-terminal Zinc fingers (residues 1-94), an adjacent segment predicted to be largely disordered (residues 95-527), and a C-terminal ATPase domain (residues 528-1167) (Fig. 2B). A point mutation predicted to inactivate the ATPase activity of TTF2^K626A^ supported normal ARP formation, CMG unloading, and CMG ubiquitylation, as did deletion of the entire ATPase domain (Fig. 2D-F; fig. S5B). In contrast, deletion of the Zn fingers inhibited these activities (Fig. 2D-F; fig. S5B). We also noticed a highly conserved, short linear motif (SLIM) comprising amino acids 120-125 that is located adjacent to the Zn fingers (Fig. 2B; blue box and fig. S5C). We therefore generated TTF2^1-200^ (Fig. 2B; red bracket), which includes the Zn finger domain and adjacent motif, and found that this construct was sufficient to restore ARP formation (Fig. 2G; fig. S5D). Therefore, the TTF2 Zn fingers and adjacent motif, whose function is documented below, are sufficient for mitotic TRAIP function, whereas, unexpectedly, the ATPase domain is dispensable.

### TTF2 interacts with TRAIP phosphorylated on T325

To understand how TTF2 promotes TRAIP-dependent CMG unloading, we used AlphaFold-Multimer (*26*) to address whether TTF2 and TRAIP are predicted to interact. AF-M detected no confident interaction between the full-length proteins (predictomes.org). Because performing structure prediction with segments of proteins increases the likelihood of detecting interactions (*27*, *28*), we divided TRAIP into individual domains and folded each with TTF2^1-200^. In this analysis, TTF2^1-200^ was predicted to interact with the C-terminal disordered region of TRAIP (fig. S6A; Table S2, rows 6 and 9), consistent with the idea that the Zn fingers contact T325. This prediction became highly confident when the proteins were further trimmed (Table S2, rows 10-11). Repeating the folding with a phosphorylated peptide in AlphaFold 3 (*29*) predicted that TRAIP phospho-threonine 325 interacts with residues K16 and R20 in a positively charged pocket between TTF2’s Zn fingers (Fig. 3A; Table S2, rows 12-13). Consistent with this prediction, TTF2^1-200^ pulled down TRAIP in a manner that was enhanced by B-CDK1, and TRAIP^T325A^ or TTF2^1-200/K16A^ did not show this effect (Fig. 3B). Underscoring the functional consequence of this interaction, TTF2^1-200/K16A^ did not support ARP formation, CMG unloading, or CMG ubiquitylation (Fig. 3C-E, and fig. S5E). Together, these results identify TTF2’s Zn finger domain as a novel phospho-peptide binding module that docks onto phosphorylated TRAIP.

**Fig. 3.**
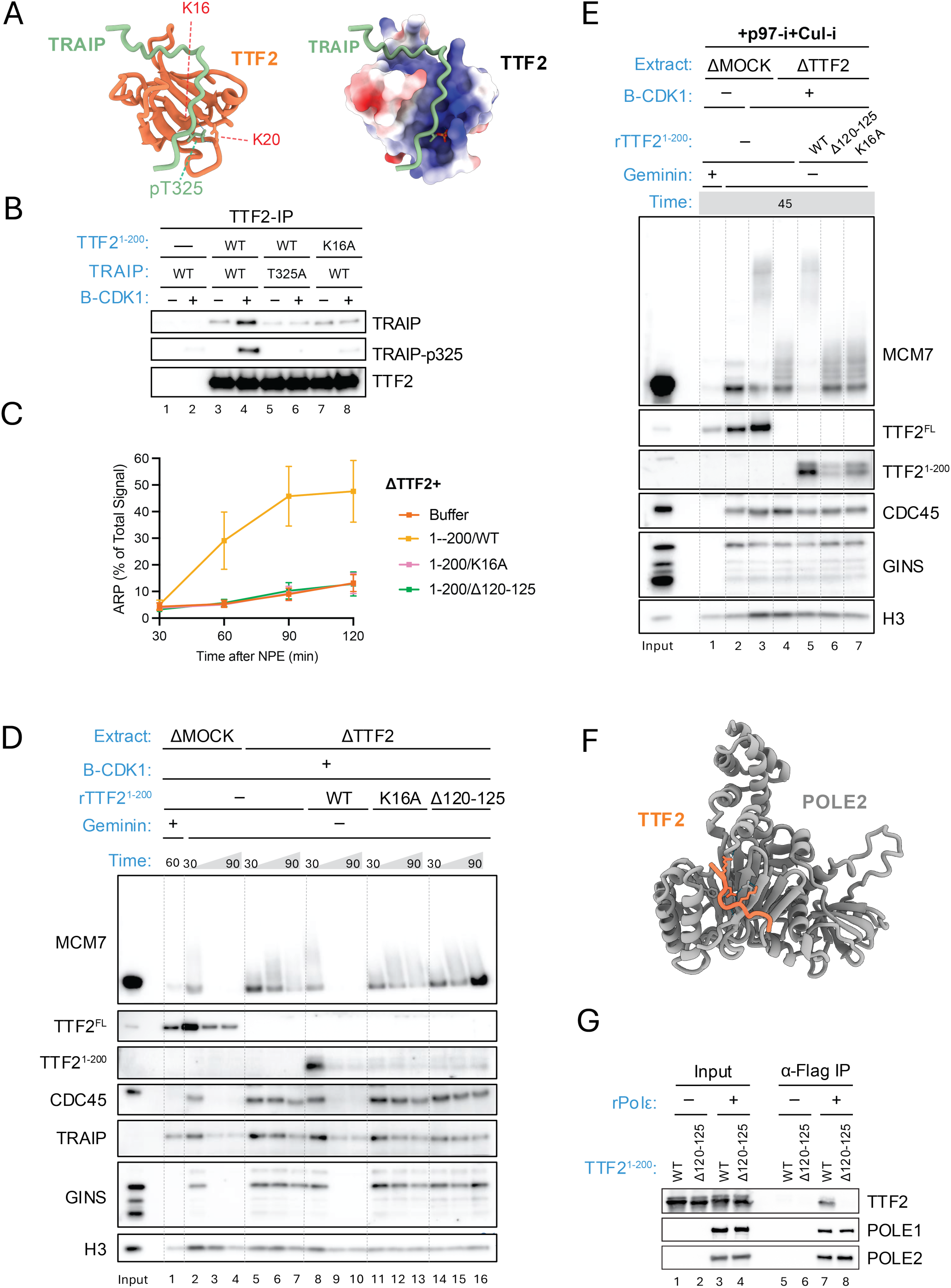
TRAIP and POLE2 binding by TTF2 is required for mitotic CMG unloading. **(A)** AlphaFold3-predicted complex between TRAIP (residues 280–339) containing phosphorylated T325 and TTF2^1–200^. Left, ribbon diagram. Right, same orientation as top with TTF2 shown with surface charge representation. **(B)** TTF2^1–200^ binds TRAIP in a phosphorylation-dependent manner. The indicated TRAIP and TTF2 proteins were expressed in TTE, B-CDK1 was added as indicated, TTF2 was recovered on beads, and pull-down samples were blotted for TTF2, TRAIP, and phosphorylated TRAIP. **(C)** The predicted interaction of TTF2 with TRAIP and POLE2 is required for ARP formation. ARP formation was quantified as in Fig. 1E in extracts supplemented with the indicated TTF2. **(D)** The predicted interaction of TTF2 with TRAIP and POLE2 is required for CMG unloading. CMG unloading was performed as in Fig. 1F using the indicated extracts and TTF2 constructs. **(E)** The predicted binding of TTF2 to TRAIP and POLE2 is required for CMG ubiquitylation. CMG ubiquitylation was performed as in Fig. 1G using the indicated extracts and TTF2 constructs. **(F)** AF-M-predicted complex of full-length TTF2 and POLE2 (see predictomes.org). Only the interacting peptide of TTF2 (118–126) is shown. TTF2 binding does not interfere with POLE2’s interaction with other replisome components. **(G)** TTF2 and pol ε interact. Anti-FLAG resin was optionally incubated with FLAG-tagged pol ε (tagged on POLE1), incubated with TTE-expressed TTF2^1–200/WT^ or TTF2^1–200/Δ120–125^, recovered, and eluted proteins were blotted as indicated.

### TTF2 also interacts with DNA pol ε

The results described above document a mitotic phosphorylation-dependent interaction between TTF2 and TRAIP, but they do not explain how this binding contributes to TRAIP function. To address this, we used a high throughput pipeline using AF-M (*30*) to fold TTF2 with ∼280 genome maintenance proteins including the 70 core DNA replication factors. Among the latter group, the most confidently predicted partner was the POLE2 subunit of DNA polymerase ε (pol ε, fig. S7A; Table S2, rows 14-15; predictomes.org) (*30*). AF-M predicted that the conserved motif adjacent to the Zn fingers (residues 120-125), which is positively charged, docks into a negatively charged groove on the surface of POLE2 that is accessible in the human replisome (Fig. 3F and fig. S5C, F). Consistent with this prediction, DNA pol ε expressed in insect cells co-immunoprecipitated TTF2^WT^ but not TTF2^1-200/Δ120-125^, which lacks the POLE2-interaction motif (Fig. 3G). Importantly, TTF2^1-200/Δ120-125^ did not support ARP formation, CMG unloading, or CMG ubiquitylation (Fig. 3C-E; fig. S5D). TTF2^1-200/Δ120-125^ also did not bind efficiently to chromatin, suggesting its function depends on being tethered to the replisome via pol ε (Fig. 3D-E). Interestingly, TTF2^1-200/K16A^, which cannot bind phosphorylated TRAIP, also bound poorly to chromatin, suggesting that TRAIP and TTF2 bind replisomes cooperatively.

### A TRAIP-TTF2 chimera bypasses the requirement for S295 and T325 phosphorylation

To further test the model that B-CDK1 phosphorylation of TRAIP promotes the interaction of TRAIP and TTF2, we generated a fusion protein in which TTF2 residues 95-205, lacking the zinc fingers but encompassing the POLE2 interaction motif, were inserted between residues 350 and 351 of TRAIP (Fig. 4A; FUSION^WT^). This chimera restored ARP formation, CMG unloading, and CMG ubiquitylation in extracts depleted of both TRAIP and TTF2 (Fig. 4B-D; fig. S8A). Importantly, the chimera still supported ARP formation even when TRAIP residues S295 and T325 were mutated to alanine (Fig. 4A-D; FUSION^2A^; fig. S8A). Thus, if TTF2 and TRAIP are covalently linked, these CDK phosphorylation sites in TRAIP are dispensable. However, when we additionally mutated the POLE2 interaction motif, the chimera failed to restore CMG unloading or ubiquitylation (Fig. 4A, C-D; FUSION^2A-ΔPOLE2^; fig. S8A). Together, these results suggest that TRAIP T325 phosphorylation allows TTF2 to bridge TRAIP’s interaction with pol ε.

**Fig. 4.**
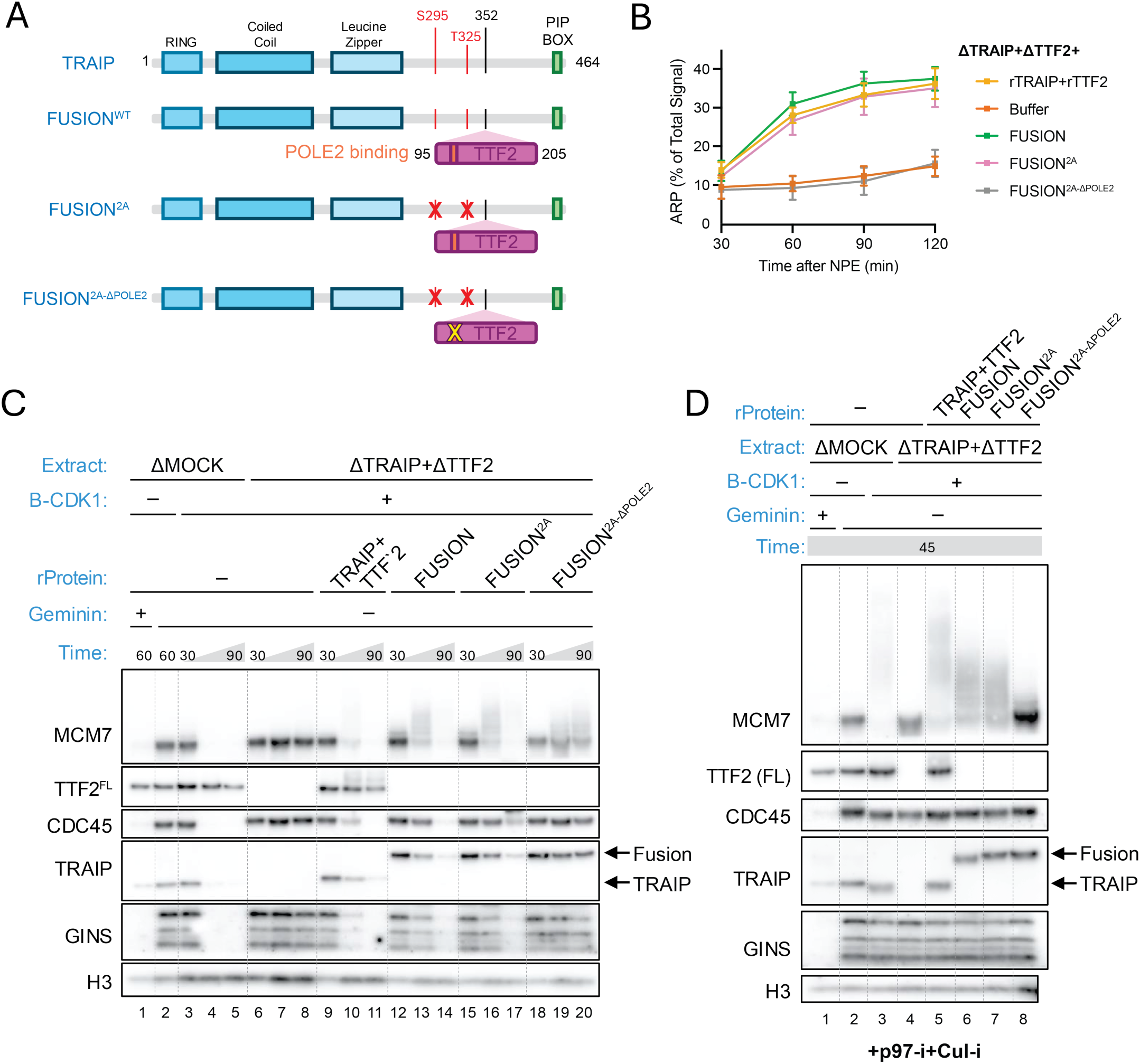
A TRAIP-TTF2 fusion bypasses the need for B-CDK1 phosphorylation. **(A)** Schematic of TRAIP–TTF2 fusions. TTF2 amino acids 95–205 were inserted between residues 350 and 351 of TRAIP. The orange box indicates the POLE2-binding region of TTF2. Red X’s indicate TRAIP S295A and T325A mutations; the yellow X indicates deletion of TTF2 residues 120–125 (Δ120–125) **(B-D)** A TRAIP-TTF2 fusion lacking CDK sites (2A), but not one also deficient in POLE2 binding (2A-ΔPOLE2) supports ARP formation (B), CMG unloading (C), and CMG ubiquitylation (D). ARP, CMG unloading, and CMG ubiquitylation assays were performed as in Figs 1E, F, and G, respectively, using mock-depleted extract or extract depleted of both TRAIP and TTF2. Depleted extract was supplemented with the indicated fusion proteins or a mixture of TRAIP and TTF2.

If the *only* function of B-CDK1 in CMG unloading is to mediate TRAIP-TTF2 binding, the TRAIP-TTF2 fusion should trigger CMG unloading in interphase. However, FUSION^WT^ did not promote ARP formation in interphase egg extract (fig. S8B, Lanes 17 to 20). One explanation is that TTF2 binding to POLE2 is blocked by DONSON, which binds to the same surface on POLE2 as TTF2 (*31*). Consistent with this possibility, DONSON appears to dissociate in mitosis (fig. S4B). Thus, TRAIP’s mitotic function requires multiple CDK1-dependent events, which we propose helps to avoid spurious CMG unloading by this pathway in interphase.

### The TRAIP-TTF2 interaction is functionally conserved

We next addressed whether the TRAIP-TTF2 axis is conserved. AlphaFold modeling showed that TTF2 is predicted with high confidence to interact with POLE2 and phosphorylated TRAIP from zebrafish to humans (fig. S6-7). Using TTE-expressed proteins, we confirmed that human TTF2^1−200^ interacts with human TRAIP in a B-CDK1-dependent manner (Fig. 5A), and this interaction depended on the hTTF2−K16 residue and the conserved hTRAIP−T324 phosphorylation site (Fig. 5A). In addition, human TTF2^1−200^ interacted with recombinant, human Pol ε complex dependent on the conserved POLE2 binding motif (Fig. 5B). Thus, the TTF2-TRAIP and TTF2-POLE2 interactions are conserved.

**Fig. 5.**
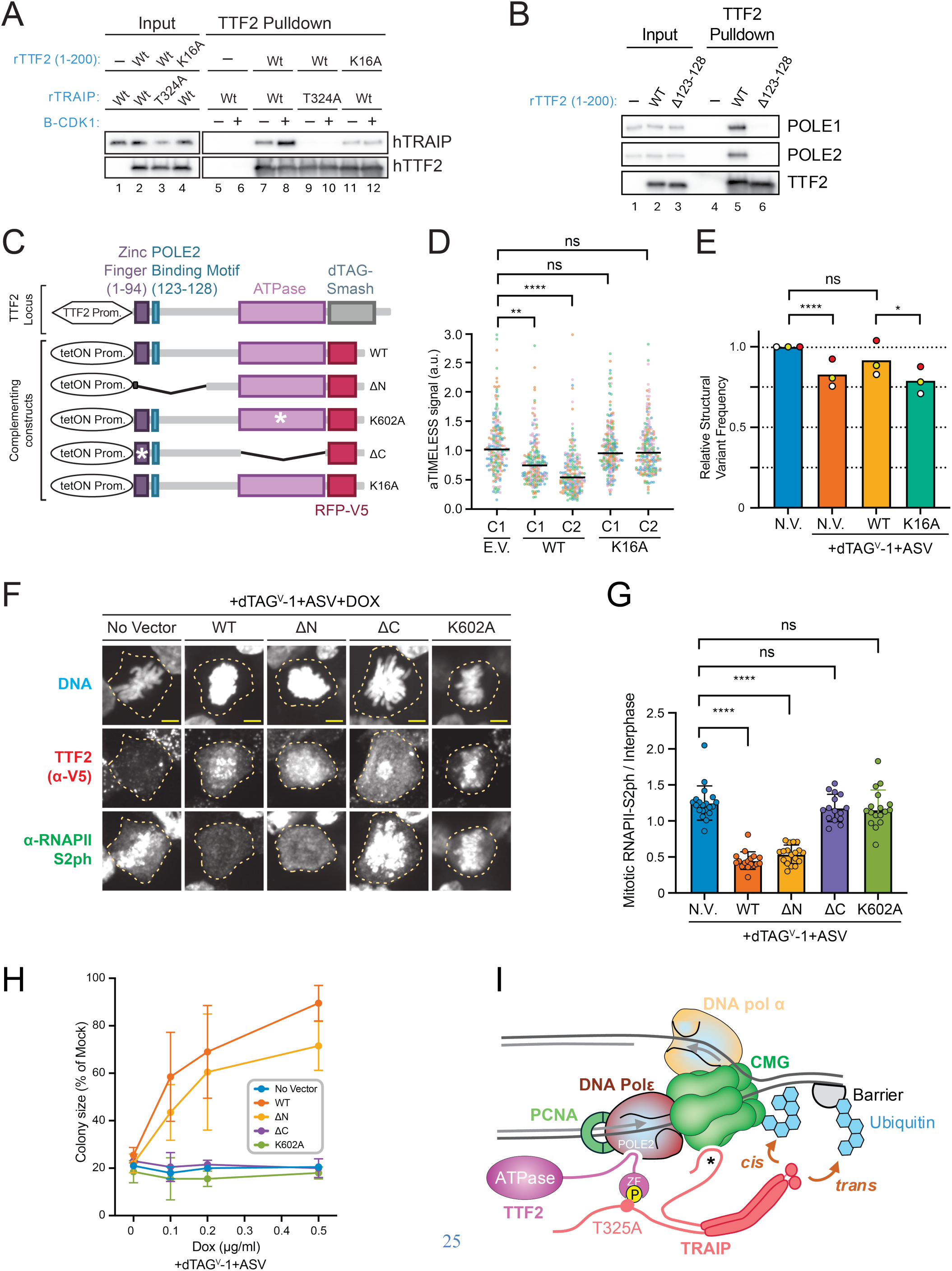
The TRAIP-TTF2 interaction is conserved in mammals and is functionally separable from ATPase-dependent RNAPII eviction. **(A)** Human TRAIP binds human TTF2. Co-immunoprecipitation was performed as in Fig. 3B except using human TTF2^1–200^ and TRAIP. Human TRAIP T324 corresponds to *Xenopus* T325. Human TTF2^1–200^ binds Pol ε. The indicated human TTF2^1–200^ constructs were incubated with purified human Pol ε, pulled down, and blotted. **(C)** Schematic of the human TTF2 locus and complementing constructs. Top, endogenous human TTF2 locus with its encoded structural domains and the engineered C-terminal dTAG-SMASh degrons. Bottom, doxycycline (DOX)-inducible complementing constructs. **(D)** TTF2 promotes replisome unloading. Endogenous TTF2 was degraded, and cells were complemented with empty vector, TTF2^WT^, or TTF2^K16A^ (one – C1, or two – C1 and C2, independent clones shown for each genotype). Cells were driven into a mitosis-like state by WEE1 and MYT1 inhibition and canonical CMG unloading was impaired by LRR1 depletion by siRNA with additional inhibition by MLN-4924. Chromatin retention of the CMG-associated protein Timeless was quantified in mitotic cells. Four experimental repeats are shown, 50 cells per sample in each repeat. Medians are shown as horizontal lines. Mann-Whitney test was used to assess the significance of differences (****, p < 0.0001). **(E)** Disrupting the TTF2-TRAIP interaction reduces deletions at CFSs. Endogenous TTF2 was optionally degraded (+dTAG^V^-1+ASV) in cells with no vector (N.V.) or in cells expressing TTF2^WT^ or TTF2^K16A^. Cells were subjected to mild replication stress, collected in mitosis, subjected to svCapture sequencing of CFS loci, and structural variants at CFS loci were quantified. Three replicates over three independent batches were normalized to the sample without TTF2 knockdown and compared using a generalized linear mixed model. **(F)** The TTF2 ATPase domain is required for RNAPII eviction. Endogenous TTF2 was degraded and TTF2 construct expression was induced (+dTAG^V^-1, +ASV, +DOX). Mitotic chromosome compaction was visualized using DNA staining, and immunofluorescence microscopy detected RNAPII (S2-phosphorylated C-terminal domain) and complementing TTF2 constructs (α-V5 tag). Cells are indicated with dashed outlines. **(G)** RNAPII-S2ph intensities from panel (F) were measured after background subtraction and normalized to interphase levels. Data are representative of three independent experiments. Fifteen cells per sample are shown. Mann-Whitney test was used to estimate significance of differences (****, p < 0.0001). **(H)** The TTF2 ATPase domain is sufficient for cell viability and proliferation. Quantification of colony size from clonogenic assays performed under endogenous TTF2 degradation and complementation with the indicated TTF2 constructs. Data are representative of four independent experiments. Mean and SD of two repeats are shown. **(I)** Model of the mitotic *Xenopus* replisome. The zinc fingers (ZF) of TTF2 bind phosphorylated TRAIP (T324 in human), and a conserved motif adjacent to the ZF binds POLE2. In this manner, TTF2 forms a flexible bridge between TRAIP and the replisome that allows *cis* ubiquitylation of CMG. The TTF2 ATPase domain is dispensable for replisome unloading but required for RNAPII eviction.

To address whether the TRAIP-TTF2 interaction promotes mitotic replisome disassembly in human cells, we modified the endogenous TTF2 locus in HCT116 colon cancer cells to encode a dTAG-SMASH tandem degron (Fig. 5C and fig. S9A), enabling TTF2 destruction within ∼30 minutes after addition of the small molecules dTAG^V^-1 and ASV (fig. S9B). We next asked whether TTF2 promotes mitotic replisome disassembly in human cells by monitoring immunofluorescence of the replisome component TIMELESS. In unperturbed cells, degradation of TTF2 did not detectably stabilize TIMELESS on mitotic chromatin, as expected given that the vast majority of replisomes are normally unloaded in S phase by CRL2^Lrr1^ (fig. S10A-C). Therefore, to assess the effect of TTF2 on active, mitotic replisomes, we depleted LRR1 and drove an asynchronous population of cells, including many in S phase, into a mitosis-like state by inhibiting the WEE1 and MYT1 kinases (*32*, *33*). In these conditions, acute degradation of TTF2 led to increased retention of TIMELESS on chromatin (Fig. 5D and fig. S10D-F). This defect was rescued by expression of wild-type TTF2 but not by the TRAIP-binding mutant TTF2^K16A^. Importantly, TTF2^K16A^ supported normal mitotic eviction of RNA polymerase II (fig. S10G-H). Therefore, the TTF2–TRAIP interaction is specifically required for efficient CMG unloading when replicating cells are driven into a mitosis-like state.

We next examined whether disruption of the TRAIP-TTF2 interaction affects genome stability. Cells were exposed to low dose aphidicolin to promote under-replication, synchronized in G2, and released into mitosis with expression of different TTF2 mutants while endogenous TTF2 was degraded. We then used svCapture sequencing to quantify structural variants at CFS loci (*34*, *35*). As predicted by our model, TTF2^K16A^ caused a decrease in the frequency of CFS deletions (Fig. 5E; fig. S11B). This effect was statistically significant, although modest (up to ∼20%). By contrast, expression of TTF2^K16A^ did not significantly affect sister chromatid exchanges, mitotic DNA synthesis, 53BP1 body formation, micronucleation (fig. S11), chromosome breakage and gap formation, CFS expression, or genomic breakage patterns detected by End-seq (figs. S12). Similar results were observed when the TTF2-POLE2 interaction was also disrupted (TTF2^K16A/Δ123−128^). Therefore, although the TTF2-TRAIP pathway promotes CMG unloading and genome maintenance in mammalian cells, its absence does not affect many genome instability endpoints, which we attribute to buffering by overlapping pathways that also process under-replicated DNA in mitosis (Discussion).

### RNAPII eviction requires the TTF2 ATPase domain and is separable from replisome unloading

Finally, we asked whether TTF2’s previously established function in RNA polymerase II eviction depends on its interaction with TRAIP. Consistent with previous siRNA results (*19*), TTF2 degradation led to a failure of RNAPII eviction from mitotic chromosomes (fig. S13A). Full length TTF2 and a mutant lacking the entire N-terminus (including the Zn Finger and adjacent POLE2 interaction site, TTF2^ΔN^) both rescued RNAPII eviction (Fig. 5F-G and fig. S13B). In contrast, a TTF2 variant with a point mutation in the Walker A motif that is expected to disrupt ATP hydrolysis (TTF2^K602A^) or TTF2 lacking the entire C-terminal ATPase domain (TTF2^ΔC^) both failed to restore RNAPII eviction (Fig. 5F-G and fig. S13B). This is consistent with prior observations that TTF2-dependent release of transcripts from stalled RNAPII *in vitro* requires ATP (*36*). Moreover, expression of TTF2^WT^ and TTF2^ΔN^, but not TTF2^K602A^ or TTF2^ΔC^ supported cell viability in cells lacking endogenous TTF2 (Fig. 5H, and fig. S13B-D). Therefore, TTF2’s essential function correlates with its ATPase-dependent transcription clearance activity. Collectively, these findings demonstrate that TTF2 promotes mitotic replisome disassembly through its N-terminal phospho-bridging function, whereas RNAPII eviction depends on its C-terminal ATPase domain.

## Discussion

Here, we identify the mechanism underlying a regulatory switch that enables a mitosis-specific response to unreplicated DNA. Previously, we showed that replisome-associated TRAIP ubiquitylates protein barriers *in trans*, whereas in mitosis it ubiquitylates the CMG *in cis*. Our finding that Cyclin B-CDK1 and TTF2 are only required for *in cis* ubiquitylation argues that TRAIP’s interphase binding mode, which remains to be elucidated, and its mitotic binding mode are fundamentally different. Mechanistically, the SLIM in TRAIP whose phosphorylation mediates TTF2 binding, and the SLIM in TTF2 that mediates POLE2 binding are both embedded in regions predicted to be disordered, which we propose confers on TRAIP the flexibility required for its catalytic RING domain to reach CMG *in cis* (Fig. 5I). Although TRAIP’s change in specificity is correlated with its structural re-organization on the replisome, we do not exclude that CDK1 also alters TRAIP’s catalytic activity, as proposed (*24*). Importantly, fusing TRAIP to TTF2 is insufficient to activate *in cis* ubiquitylation in interphase extracts. This observation strongly implies that additional CDK1-dependent events are required to allow replisome ubiquitylation and unloading, ensuring that this process never occurs during S phase, which would impede DNA replication.

TTF2 likely does not recruit TRAIP through a simple, linear pathway. We found that a TTF2 mutation predicted to disrupt TRAIP binding (K16A) also prevents efficient chromatin binding of TTF2 itself. This suggests that TRAIP and TTF2 bind replisomes cooperatively, with the two factors not only interacting with each other, but both also making independent contacts to the replisome. While TTF2 clearly contacts POLE2, a direct TRAIP-replisome contact remains to be identified. Interestingly, when we deplete TTF2, the levels of chromatin-associated TRAIP are unaffected (Fig. 3D, lanes 2 and 5). One possible explanation is that two dimers of TRAIP are associated with the replisome in mitosis, one of which is bound in the interphase mode that is not affected by loss of the functional interaction with TTF2.

An important question is whether TTF2’s function in mitotic replisome unloading is conserved in mammals. In support of this idea, we show that human TTF2 makes the same contacts as *Xenopus* TTF2 with TRAIP and pol ε, and we provide evidence that when cells are forced into mitosis, the phosphorylation-dependent TTF2-TRAIP interaction promotes replisome unloading (Fig. 3B and G; Fig. 5A, B, D, and fig. S7, fig. S10D-F). However, loss of the mitosis-specific TRAIP-TTF2-POLE2 bridge reduced SCEs and MiDAS in mES and U2OS cells (*37*) and caused a significant reduction in CFS deletions in HCT116 cells (Fig. 5E), strongly suggesting the pathway is active in cells. Nevertheless, the consequences of disrupting mitotic TRAIP function appear to depend on the biological context. Compromising the TRAIP-TTF2 interaction in mouse embryonic stem cells (*37*) or human HCT116 cells (fig. S11) had no detectable effect on classic measures of genome instability such as mitotic DNA synthesis, micronuclei, and 53BP1 body formation. Although TRAIP siRNA affects genome maintenance in U2OS cells (*17*), this might reflect its S phase function. To account for these observations, we propose that redundant mechanisms such as BLM-mediated fork dissolution can manage most of the fallout from under-replication (*38*). In the future, it will be important to address how the relative contributions of TRAIP and other pathways vary between cell types, and whether this variation represents a therapeutic vulnerability in cancer.

We showed that unlike mitotic replisome unloading, RNAPII eviction from mitotic chromosomes depends on TTF2’s ATPase activity. Moreover, the C-terminal ATPase domain is sufficient to support RNAPII eviction. Therefore, the segments of TTF2 required for replisome and RNAPII eviction are non-overlapping, arguing that TTF2 stimulates these two processes by distinct mechanisms. Whether there are circumstances, such as replisome encounter with transcription complexes in mitosis, where the two halves of TTF2 act in concert, is an interesting open question.

In conclusion, we show that CDK phosphorylation establishes a TTF2-dependent bridge that tethers TRAIP to mitotic replisomes such that it promotes ubiquitylation of CMG *in cis*. Similar results are reported in the co-submitted manuscript by Fujisawa and Labib (*37*). This structural reorganization allows cells to actively dismantle stalled replisomes before chromosome segregation, mitigating the fallout from under-replication. These findings reveal how a phosphorylation-dependent architectural switch converts an enzyme from being restrained on its host complex to attacking it, thereby repurposing the same E3 ubiquitin ligase for fundamentally different functions across the cell cycle.

## Acknowledgments

We thank Gang Zhen for assistance with the analysis of the END-seq experiments. The large language model Claude Sonnet 4.6 (Anthropic) was used for grammar and formatting assistance. All suggestions were critically evaluated, and any changes were manually incorporated by the authors.

## Funding

American Cancer Society Research Professor (J.C.W.)

National Institutes of Health grant HL098316 (J.C.W)

National Institutes of Health grant 1R35CA293978-01(D.P)

Howard Hughes Medical Institute Investigator (J.C.W, D.P)

Lustgarten Foundation for Pancreatic Cancer Research (D.P)

DFG research group FOR2800 (M.R.)

AACR-Cancer Research UK Transatlantic Fellowship (CRUK) A30783 (G.C.)

Swiss National Science Foundation (SNSF) fellowships EPM P2GEP3_191524 and PM P500PB_203132 (M.S.)

William W. Wellington Memorial Research Fund Fellowship (M.S.)

Damon Runyon post-doctoral fellowship DRF-2488-23 (A.K.)

## Author contributions

G.C. performed all experiments except those shown in Fig. 5C–D, 5F–H and figs. S9, S10, S11E–G and S13, which were performed by M.S.; figs. S4A and S11A–D, which were performed by A.K.; Fig. 5E, which was performed by S.A., with analysis by T.E.W.; fig. S12A-B, which was performed by T.W.G.; fig. S12C, which was performed by R.P. and A.N.; fig. S2B, which was performed by R.A.W.; and Fig. 3F, which was performed by Y.L. Samples for mass spectrometry were prepared by A.K. and processed and analyzed by M.R. Pol ε was generated by M.Z. G.C., M.S., J.C.W., and D.P. designed the study. G.C., J.C.W., and D.P. wrote the manuscript with input from all authors. J.C.W. and D.P. supervised the project.

## Data and materials availability

All data supporting the findings of this study, except svCapture sequencing data, END-seq data, and mass spectrometry (MS) datasets, are available in the main text or the supplementary materials. The sequencing and MS datasets below are deposited in public repositories and will all be publicly available upon publication. END-seq data have been deposited in the NCBI Gene Expression Omnibus (GEO) under accession number GSE324588. svCapture sequencing data have been deposited in the NCBI Sequence Read Archive (SRA) under BioProject accession PRJNA1403625. MS raw files and MaxQuant output tables have been deposited in the ProteomeXchange Consortium via the PRIDE repository under accession PXD055299 (CHROMASS dataset).

## Supplementary Materials

Materials and Methods

Figs. S1 to S13

Tables S1 to S3

References (*39–58*)

## Materials and Methods

### Recombinant protein expression

To purify biotinylated LacR, a plasmid expressing avidin tagged LacR pET11a[LacR-Avi] and a plasmid expressing biotin ligase pBirAcm (Avidity, Denver, CO) were co-transformed into T7 Express cells (New England Biolabs). Cultures were grown in LB media containing 50 mM biotin at 37°C (Research Organics, Cleveland, OH). Co-expression of LacR-Avi and the biotin ligase was induced by addition of 1mM IPTG (Isopropyl β-D-thiogalactoside, Sigma, St. Louis, MO). Biotinylated LacR-Avi was then purified as described (*39*), concentrated, flash frozen and stored at −80°C.

To purify DNA polymerase ε complex (pol ε), the 4 genes encoding *Xenopus laevis* POLE, POLE2, POLE3, and POLE4 were cloned individually into pLIB vectors using Gibson Assembly. POLE was cloned with a 3xFLAG tag on the N-terminus and a 10xHis tag on the C-terminus while POLE4 was cloned with a Calmodulin tag at the C-terminus. The pLIB vectors were then used to assemble a pBIG1-a baculovirus expression vector encoding all 4 subunits following the biGBac method (*40*). This vector was transformed into DH10Bac cells to generate the bacmid, and Sf9 cells were transfected with the bacmid to produce baculovirus that was amplified 3 times. The amplified baculovirus was used to transfect 1L of Sf9 cells. After 72hrs, cells were collected, washed with 1xPBS, flash frozen in liquid nitrogen and stored at −80°C. For pol ε purification, cell pellets were lysed in lysis buffer consisting of buffer H (25 mM Hepes-KOH pH=7.6, 500 mM KOAc, 10% glycerol, 1 mM DTT, 0.05% Igepal), 1 mM PMSF and 2 tablets cOmplete protease inhibitor (Roche #5056489001) per 50 mL of lysate. DNA was sheared enzymatically by benzonase and mechanically by sonication (1s ON 5s OFF for 1 min at 45% amplitude). The lysate was clarified by centrifugation at 35,000 rpm at 4°C for 1hr. Cleared lysate was allowed to bind anti-FLAG M2 affinity gel (Sigma-Aldrich #A2220-10ML) for 2hr at 4°C. The resin was equilibrated and washed extensively with lysis buffer, then the protein was eluted with buffer containing lysis buffer + 0.15mg/mL 3xFLAG peptide (Sigma-Aldrich #F4799-25MG). 2mL of Ni-NTA superflow resin (Qiagen #30430) was equilibrated with Buffer A (Buffer H + 20 mM Imidazole). Then 20 mM Imidazole was added to the eluted protein, which was then bound to the resin for 2hr at 4°C. The resin was washed extensively with Buffer A, and the protein was eluted with Buffer B (Buffer H+ 200 mM Imidazole). The eluted protein was concentrated using a 100 kDa MWCO Amicon centrifugal unit and loaded onto a superose 6 Increase 10/300 GL (Cytiva) column equilibrated with sizing buffer (Buffer H without Igepal). The protein was eluted isocratically, concentrated, flash frozen and stored at −80°C.

To purify TTF2^1-200^, the first 200 codons of *Xenopus laevis* TTF2 were cloned with N-terminal GST-3C tag into a plasmid and expressed in E.coli (GST-3C-TTF2-N). Recombinant TTF2^1-200^ protein was purified on a glutathione-sepharose column; GST was cleaved with HRV 3C protease and TTF2^1-200^ was further purified by size exclusion chromatography, concentrated, flash frozen and stored at −80°C.

Recombinant human Pol ε complex was expressed in insect cells and purified as previously described (*41*). Cyclin A2-CDK1 was purchased from Promega (Cat #V2961), Cyclin A2-CDK2 from ProQinase (Cat #0050-0054-1), and Cyclin B1-CDK1 from EMD Millipore (Cat #14-450M).

### Expression of proteins in transcription-translation extract (TTE)

TRAIP, TTF2, and FUSION (various constructs) were cloned into pF3A WG (BYDV) Flexi vectors. Mutations were introduced using a Q5 Site-Directed Mutagenesis Kit (NEB #E0554S). Plasmids were amplified in DH5α cells and purified using QIAprep Spin Miniprep Kits (Qiagen). Residual contaminants were further removed from plasmid preparations using AMPure XP Reagent (Beckman Coulter #A63881), and eluted with 10mM Tris pH 8.0. To express recombinant proteins, two volumes of 100 ng/μL TRAIP expression plasmid and 150 ng/μL of TTF2 or Fusion plasmid were mixed with three volumes of TnT® SP6 High-Yield Wheat Germ Protein Expression System (Promega # L326A) and incubated at 25°C for 2 hours. Extracts containing expressed proteins were aliquoted, flash frozen, and stored at −80°C.

### Plasmid replication substrates

The 5.5 kb LacR plasmid containing an array of 72 lacO (72xLacO) was previously described (39). A plasmid containing a site-specific cisplatin-ICL (pICLPt) was constructed as previously described (42). Briefly, parental plasmid linearized with BbsI was ligated with purified cisplatin oligonucleotide duplexes comprising Pt_Top and Pt_Bottom, and the resulting supercoiled plasmid was purified using a cesium chloride gradient.

### Xenopus egg extracts and in vitro DNA replication

Experiments involving adult female (Nasco #LM0053MX) Xenopus laevis performed at Harvard Medical School were approved by the Harvard Medical Area Standing Committee on Animals. The institution has an approved Animal Welfare Assurance (D16-00270) from the NIH Office of Laboratory Animal Welfare. Xenopus egg extracts were prepared as previously described (43).

All DNA replication reactions were performed at 22°C. To replicate pICL in interphase egg extracts (fig. S3), 1 volume of 75 ng DNA/μL plasmid DNA was added into 9 volumes of HSS and incubated for 30 minutes to promote licensing (ORC-dependent loading of MCM2-7 double hexamers). Subsequently, 1 volume HSS-DNA was mixed with 2 volumes of 50% NPE to initiate CDK2-dependent replication. To induce replication fork stalling using the LacR barrier in mitosis, one volume of 48XLacO (150 ng/µL) was incubated with one volume of 50 µM recombinant LacR for 1 hour. Next, 2 volumes of LacR-DNA were mixed with 8 volumes of HSS to allow licensing. Subsequently, 1 volume of HSS-DNA-LacR was mixed with 2 volumes of 50% NPE to initiate CDK2-dependent replication. For mitotic replication, fifteen minutes after initiation, 0.5 volumes of recombinant B-CDK1 (9 µM) was added to 14.5 volumes of the replication reaction, resulting in a final concentration of 300 nM B-CDK1. As indicated, p97 inhibitor (p97-i) NMS-873 (Sigma Cat #SML1128-5MG) and Cullin inhibitor (Cul-i) MLN-4924 (Active Biochem Cat #A-1139) were added to replication reactions 10 minutes after NPE addition at a final concentration of 250µM.

To assess CDK specificity for TRAIP phosphorylation (fig. S2E), nucleoplasmic extract (NPE) was supplemented with 500 nM Cyclin A2-CDK2, 500 nM Cyclin A2-CDK1, or 300 nM Cyclin B-CDK1 and incubated for 30 minutes at room temperature. Reactions were stopped by addition of Laemmli buffer and boiling. Endogenous TRAIP phosphorylation was detected by SDS-PAGE and immunoblotting with phospho-specific TRAIP antibodies as described above. Note that A-CDKs were used at concentrations exceeding that of B-CDK1, providing a conservative test of kinase specificity.

### Analysis of DNA replication and repair intermediates

To monitor total DNA synthesis and repair intermediates of ICL and stalled replication forks in mitosis, HSS-DNA mixtures were supplemented with 0.16 μCi/μL of [α-32P]dATP(Perkin Elmer #BLU512H500UC) just before the addition of NPE. At the indicated times after initiating replication by NPE addition, 0.8ul of the replication reactions were stopped in 9 volumes of replication stop buffer (80 mM Tris-HCl pH 8.0, 8 mM EDTA, 0.13% phosphoric acid, 10% Ficoll 400, 5% SDS, 0.2% bromophenol blue) and supplemented with 20 μg of proteinase K (Roche #3115879001). The samples were incubated at 37°C for 1 hour to digest proteins.

Samples were separated on 0.9% native agarose gels and 1X TBE buffer (89 mM Tris, 89 mM Boric acid, 2 mM EDTA pH 8.0). The gels were sandwiched between positively charged nylon membrane (Roche #11417240001) to prevent loss of nucleic acids, and paper towels to extract water, for 1 hour. Subsequently, gels were dried at 80°C under vacuum, exposed to phospho screens, and imaged on the Typhoon FLA 700 PhosphorImager (GE Healthcare).

Immunodepletions and rescue experiments

For immunodepletion of endogenous TRAIP, anti-TRAIP antibodies (Biosynth project #3472), affinity-purified as previously described (10) were used. Three volumes of the 1 mg/mL antibodies were incubated with two volumes of magnetic Protein A beads (Dynabeads M-280, Invitrogen #10001D) by gently rotating at 4°C overnight. Three volumes of 100% HSS or 55% NPE were immunodepleted by three rounds of incubation with two volumes of antibody-immobilized magnetic Protein A beads, by gently rotating at 4°C for 1 hour per round.

For immunodepletion of endogenous TTF2, a rabbit polyclonal antibody against recombinant TTF21-200 (Pocono R37741) was raised. Three volumes of 1 mg/mL affinity-purified antibody were incubated with 2 volumes of magnetic Protein A beads by gently rotating at 4°C overnight. 3 volumes of 100% HSS were immunodepleted by four rounds of incubation with 2 volumes of antibody-immobilized magnetic Protein A beads or 55% NPE was immunodepleted by three rounds of incubation with 2 volumes of antibody-immobilized magnetic Protein A beads, by gently rotating at 4°C for 1 hour per round.

For rescue experiments using TRAIP expressed in TTE, TTE was added to TRAIP-depleted NPEs at a final concentration of 6%, and the mixture was incubated for 5 minutes at room temperature before initiating replication. For rescue experiments using TTF2 expressed in TTE, TTE was added at a final concentration of 10% to NPE, and the mixture was incubated for 5 minutes at room temperature before initiating replication.

### Plasmid pull-down

Plasmid pull-downs were performed as described (44) with minor modifications. Briefly, 0.2 μM of biotinylated recombinant LacR proteins were immobilized on one volume of streptavidin-coated magnetic beads (Dynabeads M-280, Invitrogen 11206D) in six volumes of binding buffer (50 mM Tris-HCl pH 7.5, 150 mM NaCl, 1 mM EDTA, 0.02% Tween 20) for 1 hour at room temperature. The beads were washed extensively with stop buffer (20 mM HEPES-KOH pH 7.7, 50mM KCl, 5 mM MgCl2, 0.5 M sucrose, 0.25 mg/mL BSA, 0.03% Tween 20), and resuspended in six volumes of the same buffer, and chilled on ice. At the indicated time points beads were aliquoted and one volume of replication reaction sample was mixed with nine volumes of the bead aliquots and gently rotated for 30 minutes at 4°C. The beads were then washed three times with the wash buffer (20 mM HEPES-KOH pH 7.7, 50 mM KCl, 5 mM MgCl2, 0.25 mg/mL BSA, 0.3% Tween 20). Plasmid bound proteins were eluted from the beads by boiling with 1X Laemmli buffer and subjected to analysis by SDS-PAGE and immunoblotting or mass spectrometry analysis. For immunoblotting, proteins eluted from 10 ng of plasmid were loaded in each well.

### TTF2–TRAIP co-immunoprecipitation

Xenopus and human TRAIP and TTF2^1-200^ (C-terminally Twin-Strep-tagged), wild-type or mutant, were expressed separately in TTE as described above. TTE-expressed wild-type or mutant TRAIP was supplemented with 300 nM Cyclin B1–CDK1 and incubated for 30 minutes at room temperature to phosphorylate TRAIP. MagStrep® Strep-Tactin®XT beads (IBA Cat #2-5090-002) were washed three times in IP wash buffer (20 mM HEPES-KOH pH 7.7, 50 mM KCl, 5 mM MgCl₂, 500 mM sucrose, 0.25 mg/mL BSA, 0.03% Tween-20) and used in aliquots of 5 µL packed beads. For each pull-down reaction, 25 µL of TTF2^1-200^ TTE and 15 µL of TRAIP TTE (with or without phosphorylation, and with the indicated mutants) were added directly to the pre-washed beads and incubated for 1 hour at 4°C with gentle rotation. Beads were then washed three times with cold IP wash buffer. Bound proteins were eluted by two sequential elutions, each with 7.5 µL of 1× BXT buffer (IBA Cat #2-1042-025) for 15 minutes at room temperature, and the eluates were pooled. Eluates were supplemented with 5 µL of 4× Laemmli buffer, boiled, and 5 µL of each sample was loaded per lane for SDS-PAGE and immunoblotting.

### TTF2–Pol ε co-immunoprecipitation

For immunoprecipitation of Xenopus TTF2^1-200^ by Pol ε (Fig. 3G), wild-type TTF2^1-200^ and the mutant were expressed in TTE as described above. Purified recombinant FLAG-tagged Xenopus Pol ε was added at 1 µM to the TTE and incubated for 15 minutes at room temperature. Pierce™ Anti-DYKDDDDK Magnetic Agarose beads (Thermo Scientific Cat #A36797) were washed three times in IP wash buffer and used in aliquots of 0.5 µL packed beads. Each aliquot was incubated with 15 µL of the respective Pol ε–TTE mixture for 1 hour at 4°C with gentle rotation. Beads were washed three times with cold IP wash buffer. Bound proteins were eluted by incubation with 15 µL of IP wash buffer containing 1 mg/mL 3× FLAG peptide (Sigma-Aldrich #F4799) for 1 hour at room temperature with gentle rotation. Eluates were mixed with an equal volume of 2× Laemmli buffer, boiled, and 6 µL of each sample was loaded per lane for SDS-PAGE and immunoblotting.

For immunoprecipitation of human TTF2^1-200^ by human Pol ε (Fig. 5B), human wild-type TTF2^1-200^ and the mutant (C-terminally Twin-Strep-tagged) were expressed in TTE as described above. For each pull-down reaction, 25 µL of TTF2^1-200^ TTE was mixed with 5 µL of recombinant human Pol ε complex (purified as described above and diluted in IP wash buffer to a final concentration of 1 µM in the pull-down reaction) and added directly to pre-washed MagStrep® Strep-Tactin®XT beads. The mixture was incubated for 15 minutes at room temperature with gentle rotation. Beads were washed three times with cold IP wash buffer (20 mM HEPES-KOH pH 7.7, 50 mM KCl, 5 mM MgCl₂, 500 mM sucrose, 0.25 mg/mL BSA, 0.03% Tween-20). Bound proteins were eluted by two sequential elutions, each with 7.5 µL of 1× BXT buffer for 15 minutes at room temperature, and the eluates were pooled. Eluates were supplemented with 5 µL of 4× Laemmli buffer, boiled, and 5 µL of each sample was loaded per lane for SDS-PAGE and immunoblotting.

### SDS-PAGE and immunoblotting of samples from Xenopus egg extract experiments

All replication samples were quenched and boiled for 2 minutes at 95°C in Laemmli buffer (50 mM Tris-HCl pH 6.8, 2% SDS, 10% glycerol, 0.1% bromophenol blue, 5% β-mercaptoethanol). Except for Fig. 1B (see below), samples were run on 4–15% Criterion TGX Precast Midi Protein Gels (Bio-Rad #5671085) using Tris-Glycine-SDS Running Buffer (25 mM Tris-HCl pH 8.3, 192 mM glycine, 0.1 % SDS). To resolve phosphorylated TRAIP in Fig. 1B, PhosTag reagent (Wako #AAL-107) containing SDS-PAGE gels was used (*45*) with the following minor modifications. The resolving gels were composed of 6% acrylamide:bisacrylamide (29:1), 0.5% agarose, 375 mM Tris pH 8.8, 0.1% SDS, 0.001% TEMED, 30 µM ZnCl2, 0.05% APS, 30 µM Phos-tag acrylamide. The stacking gels were composed of 3% acrylamide: bisacrylamide (29:1), 125 mM Tris pH 6.8, 0.001% TEMED, 0.05% APS. Except in the case of TTF2^1-200^, gels were transferred to 0.45 µm PVDF membranes (Thermo Scientific #88518) in transfer buffer (25 mM Tris pH 8.5, 192 mM glycine, 20% methanol) at 300 mA for 1 hour. Gels for TTF2^1-200^ were transferred to 0.22 µm Nitrocellulose (GE Healthcare #10600001) membrane as described above. For Fig. 1C, the membranes were blocked with 5% (w/v) non-fat milk prepared in 1X PBST for 1 hour at room temperature by gentle agitation and incubated with primary antibodies diluted in 1XPBST containing 1% (w/v) BSA overnight at 4°C with gentle agitation. Membranes were then washed 3 times for 10 minutes with 1XPBST and incubated with secondary antibodies diluted in 5% (w/v) non-fat milk prepared in 1x PBST for 1 hour at room temperature with gentle agitation. Membranes were then washed 3 times for 10 minutes with 1XPBST and imaged using an Amersham Imager 600 (GE Healthcare). To immunoblot for phospho-specific Traip S295 and T325 antibodies in Fig. 1C, the protocol above was used except that 1X TBST and 5% BSA were used instead of 1XPBST and non-fat milk at the indicated steps. Rabbit polyclonal antibodies against phosphorylated *Xenopus* Traip S295 (Biosynth #G5978) and T325 (Biosynth #G5979) were raised by immunizing with the corresponding phosphorylated short peptides and were affinity-purified. Antibodies were further pre-cleared with B-CDK1 supplemented TRAIP depleted HSS.

For western blotting of *Xenopus* proteins, the following rabbit polyclonal antibodies were used as primary antibodies, at the specified dilutions:

TRAIP (1:10,000) (*10*)
TRAIP ps295 (1:20000, described above)
TRAIP pT325 (1:12500, described above)
MCM7 (1:12,000) (*46*)
CDC45 (1:10,000) (*46*)
GINS (1:5,000) (*47*)
TTF2 (1:2500, described above)
Histone H3 (1:500, Cell Signaling #9715, RRID: AB_331563).

The goat anti-rabbit horseradish peroxidase-conjugated secondary antibodies (Jackson ImmunoResearch, 111-035-003, RRID: AB_2313567) were used at 1:20,000 dilution for MCM7, POLE, POLE2, POLE3, POLE4 GINS and H3 blotting.

The Light chain-specific mouse anti-rabbit horseradish peroxidase-conjugated secondary antibodies (Jackson ImmunoResearch, 211-032-171, RRID: AB_2339149) were used at 1:10,000 dilution for TRAIP, TTF2, and CDC45 blotting.

### MS sample preparation

For plasmid pull down, 8 µl of the total reaction was withdrawn at the indicated time points and plasmids and associated proteins were recovered by plasmid pull down using LacI coated beads as described above with minor modifications. The samples were washed twice with 20 mM HEPES-KOH (pH 7.7), 100 mM KCl, 5 mM MgCl2, 0.25 mg ml−1 BSA, and 0.03% Tween-20 and once with 50 mL ELB (10 mM HEPES/KOH at pH7.7, 50 mM KCl, 2.5 mM MgCl2) to remove residual detergent. The beads were diluted with ABC (50 mM ammoniumbicarbonate) and digested with 2.5 mg trypsin (Sigma) and 2M Urea and incubated for 16 hours at 30 °C. Supernatant was cleared by spinning through 0.45μm ultrafree filters before stopping the digestion with Trifluoroacetic acid. NaCl was added to 400 mM final concentration and peptides were acidified and purified by stage tipping on C18 material (48).

### LC-MS/MS analysis

Peptides were separated on reversed phase columns (50 cm, 75 μm inner diameter, packed in-house with ReproSil-Pur C18-AQ 1.8 μm resin (Dr. Maisch GmbH) and directly injected into a quadrupol orbitrap mass spectrometer (Q Exactive HF, Thermo Scientific, Germany). Using a nanoflow HPLC (Thermo Scientific, Odense), peptides were loaded in buffer A (0.5% acetic acid) and eluted with a three hour non-linear gradient from 5-95% buffer B (80% acetonitrile, 0.5% acetic acid) at a constant flow rate of 250 nl/min. For the reversed phase separation, the column was maintained at a constant temperature of 60°C. The mass spectrometer was operated in a data dependent fashion using a top 15 method for peptide sequencing.

### MS Data processing

Raw data were analyzed with the MaxQuant software (version 2.0.1.0) (49). A false discovery rate (FDR) of 0.01 for proteins and peptides and a minimum peptide length of 7 amino acids were required. MS/MS spectra were searched against a non-redundant Xenopus database (see (50) for details). For the Andromeda search, trypsin allowing for cleavage N-terminal to proline was chosen as enzyme specificity. Cysteine carbamidomethylation was selected as a fixed modification, while protein N-terminal acetylation and methionine oxidation were selected as variable modifications. Maximally two missed cleavages were allowed. Protein identification required one unique peptide to the protein group. Raw intensities were normalized using the label free quantification (LFQ) algorithm implemented in MaxQuant. Match between run was enabled to transfer identities between runs of the same replicate group.

### Statistical analysis and visualization of MS data

Protein groups were filtered to eliminate contaminants, reverse hits, and proteins identified by site only. For the heat map (Fig. 2A and fig. S4B), LFQ intensities were log2 transformed and for each protein z-scores were calculated across all conditions. To visualize heatmaps in a more compressed form, z-scores of individual subunits were averaged for selected complexes (see Table S1 for the z-scores of all proteins). Proteins were manually annotated and sorted according to their function in DNA replication and DNA repair. To identify proteins with significant abundance changes between the four conditions, LFQ intensities were log2 transformed, and missing values were imputed with random values drawn from a normal distribution centered around the detection limit of the MS instrument (Perseus imputation; width=0.3, down shift = 1.8). Two sample Student’s t-tests with permutation-based FDR control were carried out in Perseus. For these tests, three valid values in at least one triplicate of either of the tested conditions was required. FDR was adjusted for multiple testing by the Benjamini-Hochberg procedure using a significance threshold of FDR<0.05 (see Table S1). Data visualization was carried out in R. All scripts are available upon request.

### Cell lines and Constructs

The HCT116 cell line and its derivatives were grown in McCoy’s 5A medium (30-2007, ATCC) supplemented with 10% tetracycline-free FBS and 1% penicillin-streptomycin mixture in 5% CO2 at 37°C. TTF2-degron cell lines were created using CRISPR Cas9 genome editing with marker-free co-selection (51). To this end, HCT116 cells were transfected using Neon Transfection System with a mixture of three DNA vectors: (1) double Cas9-gRNA vector (containing Cas9 and gRNA against TTF2 STOP codon and ATP1A1 ouabain suppression mutation site; eSpCas9(1.1)_gATP1A1_gT2C2), (2) TTF2 degron HDR targeting template (hTTF2_CT_dTAG_SMASh2), and (3) ATP1A1 ouabain resistance HDR targeting template (ATP1A1_Q11RN129D). Transfected cells were selected for growth in ouabain (10 μM; O3125, Sigma) containing media. Resistant cells were FACS sorted to single cells in 96-well plates, and clones were screened by PCR for degron cassette integration and by immunoblotting for TTF2 depletion under degron activation (Table S3). TTF2-degron depletion was induced by supplementing media with 0.5 μM dTAG^V^-1 (Cat #6914, Tocris)(52) and 1 μM ASV (Cat #HY-14434, MedChemExpress)(53).

To complement the TTF2-degron cell lines with ectopically expressed doxycycline (DOX) inducible TTF2 versions, the human TTF2 open reading frame was cloned into a pLIX403 lentiviral vector under tetON promoter with C-terminal V5 affinity tag and RFP fluorescent protein tag. The resulting pLIX403_hTTF2-V5-RFP was mutagenized by PCR and NEBuilder HiFi assembly kit cloning to obtain TTF2 mutant constructs. pLIX403 based vectors were then used to produce lentivirus. Lentivirus-infected TTF2-degron cell lines were selected on puromycin, FACS sorted for RFP expression (under transient DOX treatment), and single-cell cloned to obtain clonal cell lines. TTF2 ectopic expression was induced by addition of 0.1-0.5 μg/mL DOX in the media for 24 hours or longer. All cell lines tested negative for mycoplasma contamination.

### Immunofluorescence and Microscopy

#### RNA polymerase II staining in fixed cells

Cells were seeded on poly-D-lysine coverslips in 12-well plates at a concentration of 50 k/mL (day 1). On day 4, the coverslips with cells were washed with PBS and fixed with simultaneous permeabilization using PTEMF solution (20 mM PIPES pH 6.8, 1 mM MgCl2, 10 mM EGTA, 0.2 % Triton X-100, 4 % paraformaldehyde) at RT for 15 min. Coverslips were washed thrice with PBS (5 min each wash here and after) and kept at 4 °C until staining (up to a week). For immunostaining, fixed coverslips were blocked with blocking buffer (3 % BSA in PBS) for 1 hour at RT, incubated with primary antibodies diluted in blocking buffer for 1 hour at RT, washed thrice with Wash buffer (0.05 % Triton X-100 in PBS), and incubated for 1 hour at RT with secondary antibodies diluted in blocking buffer. After removing secondary antibodies and three more washes with Wash buffer, the coverslips were incubated with 1:2500 dilution of Hoechst stain (H3570, Invitrogen) for 10 min at RT. Hoechst solution was removed, and coverslips were washed thrice with PBS. Coverslips were mounted in Prolong Diamond Antifade (P36965, Invitrogen) on imaging slides, cured overnight at RT and imaged using a spinning disc confocal microscope (Nikon) with a 60X objective. Images were processed in Fiji. Z-stacks were max intensity projected. Primary antibodies: α-RNAPII-S2ph 1:400 (ab5095, rabbit, Abcam), α-V5 1:200 (R960-25, mouse, Thermo), α-TTF2 1:100 (sc-514996, mouse, SantaCruz BioTechnology). Secondary antibodies: goat anti-rabbit Alexa Fluor 488 1:1000 (A32731, Invitrogen), goat anti-mouse Alexa Fluor 568 1:1000 (A11031, Invitrogen).

#### TIMELESS and RNA polymerase II staining on mitotic chromatin

For TIMELESS and RNAPII detection on mitotic chromatin, cells were seeded directly in 12-well plates (day 1). On day 2, Doxycycline (DOX, 1 μg/mL) was added to induce ectopic TTF2 or empty vector expression. On days 3 and 4, cells were transfected with siRNA against dTAG portion of the endogenous TTF2-degron (ON-TARGETplus Human FKBP1A (2280) siRNA – SMARTpool, L-009494-00-0005, Horizondiscovery/Dharmacon) and against LRR1 (ON-TARGETplus Human LRR1 (122769) siRNA – SMARTpool, L-016820-01-0005, Horizondiscovery/Dharmacon), 40 nM each. Briefly, per one well of a 12-well plate, 50 uL of Opti-MEM (31-985-062, Gibco) was mixed with 4 uL of 10 μM siRNA stock solution. In parallel, 50 uL of Opti-MEM was mixed with 2.5 uL of Lipofectamine 3000 (L3000015, Invitrogen). Both tubes were combined, mixed, and incubated 5 min at RT. The media in wells was refreshed (while also replenishing DOX) and the transfection mixture applied into wells dropwise. The plates were gently swirled and returned to incubator. On day 4 (18 hours before cell collection) MLN4924 (2 μM final, A-1139, Active Biochem) was added to inhibit CRL2^Lrr1^ and dTAG^V^-1 (0.5 μM) and ASV (1μM) were added to induce TTF2-degron degradation. Sixteen hours after, WEE1 (10 μM, AZD1775, S1525, Selleckchem) and MYT1 (10 μM, lunresertib, HY-145817A, MedChem Express) kinase inhibitors were added to induce mitotic entry. Two hours after, floating cells from each well were collected in tubes (1 mL) and spun down 500 g x 5 min at RT. All but 150 uL of supernatant was discarded and the cell pellet was re-suspended in remaining supernatant. Concentrated cells were split in three equal fractions: two applied onto poly-D-lysine coverslips in new 12-well plates (for aTIMELESS and aRNAPII staining) and one was used to prepare protein lysates for polyacrylamide gel electrophoresis and Western Blotting. The 12-well plates with coverslips and cells on them were returned to 37C incubator for 10 minutes to let cells attach.

To detect chromatin-bound proteins, we followed a protocol previously published by Meyer Laboratory (54). Briefly, the coverslips were washed once with PBS, pre-extracted 5 minutes on ice (500uL/well ice-cold 0.2% Triton-X100 (TX100, AAA16046AP, FisherScientific) with 1x cOmplete protease inhibitor (5056489001, Roche) and then fixed by adding equal volume of 8% PFA in PBS (4% PFA final in the well) followed by 15-minute incubation at RT. The coverslips were then washed three times with PBS and stored at 4C up to one week. To stain, cells were permeabilized again (1000uL/well 0.2% TX100 (in PBS), 15 min at RT) and blocked with 1 mL/well of blocking buffer II (10% FBS, 1% BSA, 0.1% TX100 in PBS) for 1 hour at RT. The coverslips were then incubated with primary antibodies diluted in blocking buffer II for 2 hours at RT, washed three times with PBS, incubated with secondary antibodies diluted in blocking buffer II for 1 hour at RT. The secondary antibody solution was aspirated, and DNA stain (1:2500 dilution of Hoechst, H3570, Invitrogen) was applied onto coverslips for 10 min at RT. The coverslips were then washed three times with PBS and mounted onto glass slides as above for RNA polymerase II staining in fixed cells. Microscopy imaging of coverslips from the same experimental replicate were done on the same day using identical imaging settings.

Primary antibodies: Anti-Timeless antibody [EPR5275] 1:800 (ab109512, rabbit, Abcam), α-V5 1:200 (R960-25, mouse, Thermo), anti-CENPC 1:5000 (PD030MS, guinea pig, mblintl)

Secondary antibodies: goat anti-rabbit Alexa Fluor 488 1:1000 (A32731, Invitrogen), goat anti-mouse Alexa Fluor 568 1:1000 (A11031, Invitrogen), Goat anti-Guinea Pig Alexa Fluor 647 1:1000 (A21450, Invitrogen).

#### 53BP1 foci and micronuclei staining

For 53BP1 foci and micronuclei imaging, cells were seeded on coverslips in 12-well plates as above and allowed to attach overnight. Doxycycline (DOX, 1 μg/mL) was added on the next day to induce ectopic TTF2 expression. Aphidicolin (APH, 0.4 μM, A0781-5MG, Sigma) was added 38 hours after DOX, and RO-3306 (9 μM, SML0569-5MG, Sigma) was added 11 hours after aphidicolin. At two hours before release, cells were treated with dTAG^V^-1 (0.5 μM) and ASV (1 μM) to degrade endogenous TTF2-degron. After 13 hours of incubation with RO-3306, the cells were released in mitosis by four washes with warm media (supplemented with DOX, dTAG^V^-1 & ASV). Cells were then washed with PBS, fixed with 4% paraformaldehyde at RT for 15 min, washed thrice with PBS and stored at 4 °C until staining (up to a week). Cells on coverslips were permeabilized with 0.5% Triton X-100 in PBS for 5 min at RT, washed three times with PBS. Blocking, staining and imaging was done as described above for RNA polymerase II staining in fixed cells.

Primary antibodies: Purified α-53BP1 1:300 (clone W17184B, rat, 933001, BioLegend), α-V5 1:200 (R960-25, mouse, Thermo), α-Cyclin A2 (E6D1J) 1:1000 (67955T, rabbit, Cell Signaling Technology)

Secondary antibodies: goat anti-rat Alexa Fluor 488 1:1000 (A11006, Invitrogen), goat anti-mouse Alexa Fluor 568 1:1000 (A11031, Invitrogen), goat anti-rabbit Alexa Fluor 647 1:1000 (A21245, Invitrogen).

### Clonogenic proliferation assay

Cells were seeded in 6-well plates at 200, 400, 600 (fig. S13C), or 450 cells per well (Fig. 5H, fig. S13D) (day 1). Where indicated, doxycycline (DOX) was added on day 2 and degron ligands (dTAG^V^-1 and ASV) were added on day 3. Medium and drugs were refreshed every 3 days. On day 14, media was aspirated, the wells were washed with PBS, colonies were fixed by methanol for 20 min at RT and stained by crystal violet solution (0.4 % (w/v) crystal violet, 20 % methanol) for 1 hour at RT. The crystal violet solution was then recovered, and wells washed thrice with warm tap water. The plates were dried overnight and photographed with digital camera; images were cropped in ImageJ.

### Clonogenic assay image quantification

Clonogenic assay plate images were analyzed in ImageJ and Excel. Each plate image was converted to 8-bit grayscale (Image > Type > 8-bit), and thresholded by limiting contrast to 0-150 (Image > Adjust > Threshold 0-150). To separate colonies, the watershed method was then applied (Process > Binary > Watershed). Wells were selected with the circle select tool of equal size per plate, and colonies were detected and measured using the Analyze Particles tool. Average colony size in percents was calculated in Excel by dividing the average colony size in each well by that of the mock-treated well and multiplication by 100. Average colony size values were then reported on the figure graph using Prism.

### Immunofluorescence microscopy image quantification

#### Image analysis of RNA polymerase II staining in fixed cells

The immunofluorescence microscopy images were analyzed using custom Fiji macro and Excel. In Fiji, image hyperstacks were first collapsed (Image > Stacks > Z project > MAX intensity option), and individual channels set to grays (LUT > Grays). To measure signal intensity for each channel on and outside of chromatin, a nuclear (DNA) mask was first created. To this end, the DNA channel (Hoechst) image was duplicated, converted to 8-bit (Image > Type > 8-bit), smoothed by applying Median filter (with 3 pixels setting) and thresholded with ‘Huang’ method (Image > Adjust > Threshold (Huang)). Holes were filled (Process > Binary > Fill Holes), chromosome groups were separated using watershed (Process > Binary > Watershed) and obtained DNA mask was saved. Chromosome groups (interphase nuclei and mitotic chromosomes) were detected on the mask by using Analyze Particles method (with size set to 1000-Infinity). Each channel signal intensity was then measured for all chromatin groups Regions of Interest detected with the mask (ROI Manager > measure). To obtain the signal intensity outside of chromatin (background), the DNA mask was first inverted (Edit > Invert), total region outside of chromatin selected (Edit > Selection > Create Selection) and used for quantification in ROI Manager (Analyze > Tools > ROI Manager + Add > measure). Obtained measurements were imported to Excel, where RNAPII-S2ph background intensity was subtracted from intensity on chromatin. The RNAPII-S2ph signal intensity on TTF2-positive (or negative, for no_vector control) mitotic chromatin was divided by average signal on interphase nuclei of the same field of view, and obtained values were reported on the figure graph using Prism. Prism Mann-Whitney U test was used to test for statistical significance of the difference in value distributions.

#### Image analysis of TIMELESS and RNAPII on mitotic chromatin

The same Fiji macro as for RNA polymerase II staining in fixed cells was used. A custom Python script was then applied to subtract background signal intensity for each field of view, filter cells from debris by size (Area between 4000 and 20’000) and select mitotic cells (by thresholding CENPC signal Skew > 3). The data was then imported to Excel. For samples with ectopic TTF2 expression, additional filter was applied to select only TTF2-positive nuclei (V5 signal > 40). The above background per nucleus mean intensities of TIMELESS and RNAPII staining were then normalized to the median of ‘empty vector’ sample values distribution from the same experiment and same image acquisition day. Prism Mann-Whitney U test was used to test for statistical significance of the difference in value distributions.

### Immunoblotting of cell-based samples

Cells were trypsinized, pelleted, washed with PBS, re-suspended in 2x Laemmli buffer and boiled for 5 minutes at 95°C. Denatured samples were resolved on 4-15% Tris-Glycine SDS PAGE, transferred to PVDF membrane, blocked with milk, washed thrice with PBS, and incubated with primary antibody overnight at 4°C with shaking. Primary antibody was decanted, membrane was washed thrice with PBS and incubated with secondary antibody for 45 minutes at RT with shaking. After three washes, the membrane was incubated with SuperSignal ECL (Thermo) and imaged with Amersham Imager. Images were cropped in ImageJ. Primary antibodies: α-TTF2 1:5,000 (PA5-96789, rabbit, Invitrogen), α-HA 1:1,000 (C29F4, rabbit, Cell Signaling), α-RPS19 1:1,000 (A304-002A, rabbit, Bethyl), α-V5 1:5,000 (R960-25, mouse, Thermo). Secondary antibodies: goat anti-rabbit HRP 1:30,000, rabbit anti-mouse HRP 1:2,000.

### Cell culture and treatment for common fragile site (CFS) analysis and svCapture

Cell growth and treatment for common fragile site (CFS) analysis and svCapture were primarily the same with a few minor differences. HCT116 (ATCC, RRID CVCL_0291) cells were maintained at 37 °C in a humidified incubator with 5% CO₂ using McCoy’s 5A (modified) medium (Gibco, 16600082) supplemented with 10% fetal bovine serum, 2 mM L-glutamine, and 100 U/mL penicillin–streptomycin. Cells were treated with 0.5 μg/mL doxycycline to induce TTF2 plasmid expression for 16 hours followed by addition of 0.4 µM aphidicolin for 6 hours and then arrested at G2/M with 9 µM RO3306 for an additional 18 hours. Cells were washed three times with PBS and released into warm media containing 75 ng/ml Colcemid (Gibco) for 30-45 minutes for CFS analysis or 3 hours for flow sorting of G2 and M fractions. Asunaprevir (1 µM) and dTAG^V^-1 (0.5 µM) were added 2 hours prior to RO3306 release and again following release into fresh media.

### Common fragile site analysis

After harvesting cells with trypsinization, chromosome preparations were obtained by a 15-minute incubation in 0.075 M KCl hypotonic solution at 37 °C followed by multiple washes with Carnoy’s fixative (3:1 methanol:acetic acid). Fixed cells were dropped onto glass slides to produce metaphase spreads, which were then baked overnight at 60°C prior to Giemsa banding. For banding, slides were rinsed in water, digested with trypsin solution (0.0005% trypsin and 0.02% Tyrode’s in HBSS) for 50 seconds, washed twice in 0.9% NaCl, and stained for 5 minutes with Giemsa solution (5% Giemsa in Gurr buffer, pH 6.8), followed by two final water rinses. G-banded metaphase chromosomes were examined using a Zeiss Axiphot microscope. Gaps and breaks specifically at the CFSs FRA3B (3p14) and FRA16D (16q23) and total chromosomal gaps and breaks were scored in 25 metaphases per experimental condition.

### svCapture analysis

Trypsinized cells were collected in cold media, centrifuged for 5 minutes at 500×g at 4 °C, and fixed overnight at −20 °C in 70% ethanol at a density of 1×10⁶–2×10⁶ cells/mL. Fixed cells were washed with PBS, permeabilized on ice for 15 minutes using 0.25% Triton X-100 in PBS and centrifuged. Cells were then incubated for 1 hour with an Alexa Fluor 488–conjugated phospho-histone H3 (pH3) antibody (Cell Signaling Technology; Invitrogen) diluted 1:50 in antibody staining buffer (0.5% BSA in PBS). After two washes with staining buffer, cells were stained with propidium iodide (100 µg/mL) and RNase (100 µg/mL). Samples were analyzed at the University of Michigan Flow Cytometry Core using Bigfoot Cell Sorter (Thermo Fisher). Cell cycle gating defined G2-phase cells as 4N DNA content and pH3 negative, and M-phase cells as 4N DNA content and pH3 positive. At least 200,000 cells were collected across all targeted cell cycle fractions.

Flow-sorted samples were centrifuged at 1,000 rpm for 10 minutes. The supernatant was removed, leaving 100 µl, to which 200 µl of DNA/RNA Shield (Zymo Research, R1100-50) and 15 µl of proteinase K (20 mg/ml) were added. Samples were incubated at room temperature for 20 minutes. Genomic DNA was then isolated using the Quick-DNA Microprep Plus kit following the manufacturer’s protocol (Zymo Research). Sequencing libraries were generated at the University of Michigan Advanced Genomics Core using the Illumina DNA Prep with Enrichment kit with 300 ng of genomic DNA, IDT for Illumina unique dual indexes and amplified by nine PCR cycles. Library concentration was measured using Qubit, and fragment size and quality were assessed with an Agilent TapeStation to confirm 350 ng-500ng of library material. Hybridization capture probes targeting the central 400 kb of the WWOX and FHIT genes were designed and synthesized by Twist Biosciences. For capture, 500 ng of each library was pooled and hybridized with 4 µl of Twist probes and 6 µl of PCR-grade water. Target enrichment was performed using magnetic beads according to Illumina kit guidelines, followed by amplification of captured fragments with 12 PCR cycles. Sequencing used paired-end 2 x150 bp reads on either a Illumina NovaSeq X Plus or AVITI 24 sequencer.

svCapture data analysis was performed using the pipeline of the same name from the svx-mdi-tools code repository available on GitHub (https://github.com/wilsontelab/svx-mdi-tools) with releases mirrored on Zenodo (https://doi.org/10.5281/zenodo.7871676). A complete description of the pipeline actions has been provided (*34*, *35*), which entailed read alignment to the human hg38 reference genome, read de-duplication, SV junction detection, characterization, and unique molecule counting, and coverage assessment. The initial result from each sample, SV Frequency, was calculated as the number of unique deletion SVs from 10 kb to 1 Mb detected only once in one sample (the expectation for newly formed SVs) divided by the average read coverage in the capture target regions (a measure of data depth per sample). Due to observed batch effects between sample sets, we further normalized the SV Frequency in each co-analyzed batch of samples to the value from the APH-induced M-phase sample with TTF2 expression from the native gene (expected to have the highest SV yield) from that batch. Statistical differences between sample groups were assessed using a generalized linear model based on the negative binomial distribution that accounted for sample read depth as well as the experimental groups and batches as independent co-variates, as previously described (*34*).

### Mitotic DNA synthesis (MiDAS)

Cells were seeded onto 10 cm dishes and induced with 1 μg/ml doxycycline the next day (Day 2). On day 3, cells were treated with 0.4μM aphidicolin (Sigma) for 16h. The following day (Day 4), cells were arrested in G2 phase of cell cycle by treatment with 9 μM CDK1 inhibitor (RO-3306, Selleck) for 6h. 2h before release, cells were treated with dTAG^V^-1 and ASV to degrade endogenous TTF2. To release cells from G2 arrest, cells were washed in warm HBSS three times and released in fresh medium with 20 μM EdU supplemented with dTAG^V^-1 and ASV for 25 min. Subsequently, mitotic cells were collected in conditioned media with shake off. The cells were pelleted by centrifugation at 900xg for 5min and the pellet was resuspended in 250ul media and plated on poly-lysine coated slides. Slides were fixed with PTEMF buffer (4% paraformaldehyde (PFA),200 mM HEPES pH 6.8, 200 mM MgCl2, 10 mM EGTA pH 8.0, 0.2% Triton X-100) at room temperature for 10min and washed thrice with PBS before storing the slides at −20 °C until staining. EdU click reaction was performed per the manufacturer’s instructions by incubating the slides in dark for 1h. The slides were blocked with 3%BSA/PBS with 0.1% Triton for 1h before incubating in blocking solution overnight. The following day, slides were rinsed three times with PBS and mounted with Prolong gold DAPI (Life technologies). All images were acquired on Nikon CSU-W1 SoRa with 63x oil objective. The number of foci per mitotic cell were blinded and scored manually. For each experiment, at least 100 mitotic cells were scored.

### Sister Chromatid Exchange (SCE)

SCE was performed essentially as previously described with a few modifications (*55*). Briefly, cells were seeded on 10cm dishes and 10μM BrdU (Life Technologies) and 1 μg/ml Dox were added the following day. The day after that, 0.4μM aphidicolin was added and 6h later 9μM RO-3306 was added for 16h to arrest cells in the G2 phase. 2h before release from G2, cells were treated with dTAG^V^-1 and ASV to degrade endogenous TTF2. To release cells into mitosis, the cells were washed with warm HBSS and released into fresh medium containing dTAG^V^-1, ASV, and 0.075 μg/ml colcemid (Roche) for 45 minutes. Cells were harvested in conditioned media by trypsinization and fixed in cold methanol: acetic acid (3:1). Metaphase spreads were prepared and stained the next day. Slides were immersed in 10 μg/ml Hoechst 33258 (Life technologies) for 20 min in the dark at RT and exposed to UV-A for 60 minutes. Subsequently, slides were incubated in 1X SSC (Invitrogen) at 60 °C for 1h before rinsing in distilled water three times. The slides were then mounted with Prolong gold DAPI and images were acquired on Nikon CSU-W1 SoRa with 100x oil objective. Slides were blinded and SCEs were scored manually. At least 80 cells were scored per condition.

### END-Seq analysis

END-seq was performed as previously described (56). Briefly, HCT116 cells were embedded in agarose plugs and treated with Proteinase K and RNase A. Adapter ligation, DNA sonication, and library preparation were performed as previously described (57).

### Use of AI

We asked the large language model Claude (Sonnet 4.6 version) for feedback on our manuscript. All changes implemented were fully vetted and manually incorporated into the text by the authors with further editing.

---

**Fig. S1.**
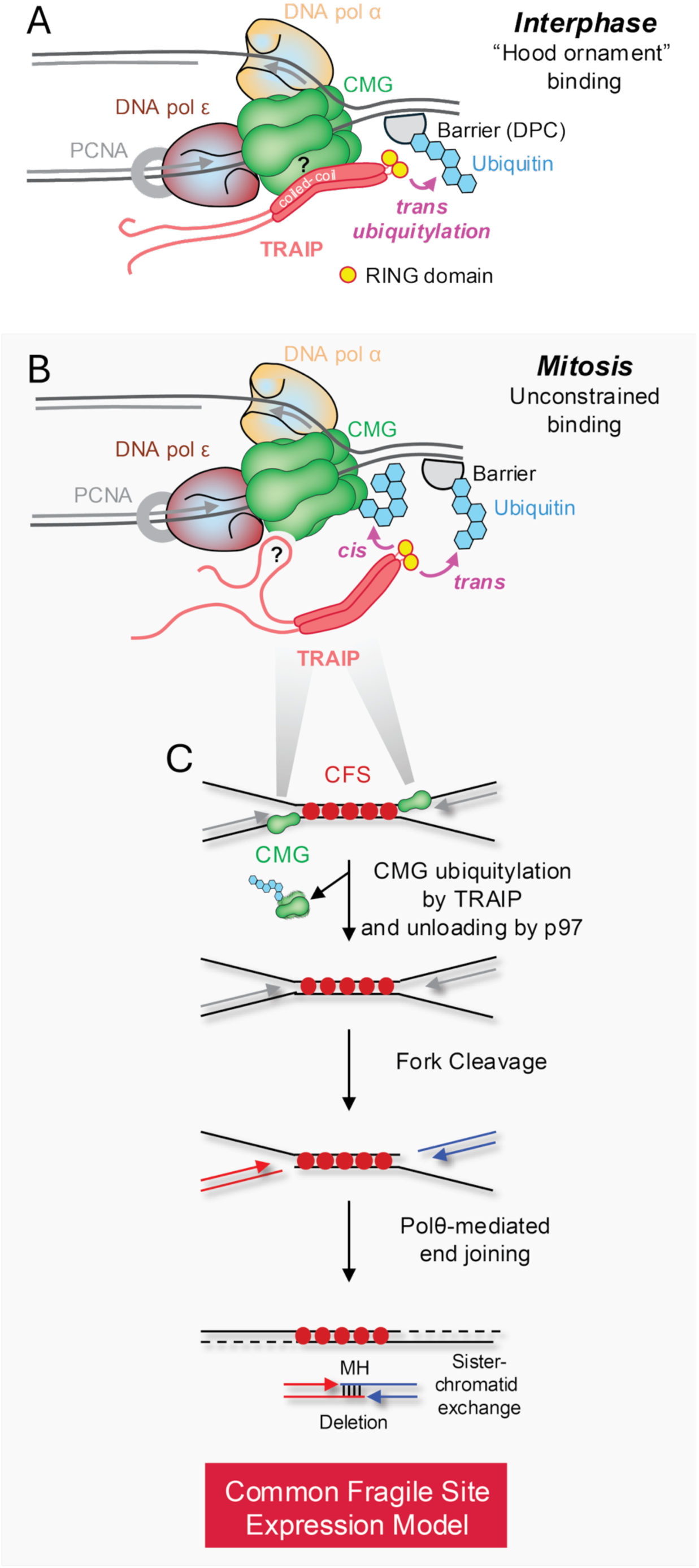
Models for TRAIP replisome binding in the S and M phases. **(A)** Model for the interphase configuration of TRAIP on the replisome. The TRAIP dimer binds the replisome such that the catalytic RING domains (yellow circles) ubiquitylate proteins ahead of the replisome in *trans* (“Hood ornament” model). In this configuration, TRAIP ubiquitylates proteinaceous barriers ahead of the fork such as covalent DNA protein cross-links (DPCs) or a converged CMG at a DNA interstrand cross-link (not depicted), but it cannot ubiquitylate the CMG with which it travels *in cis*. **(B)** Model for the mitotic configuration. TRAIP is attached to the replisome such that it can ubiquitylate the host CMG in *cis*. **(C)** Model for the processing of unreplicated DNA at common fragile sites (CFS)(*14*). If replication does not complete before cells enter mitosis, the stalled CMGs undergo TRAIP-dependent ubiquitylation. The resulting p97-dependent replisome unloading deprotects the forks and induces symmetric fork cleavage. The broken sister chromatids (red and blue) are ligated by pol θ-mediated end-joining, leading to a small deletion with microhomology (MH) at the breakpoints and sister chromatid exchange.

**Fig. S2.**
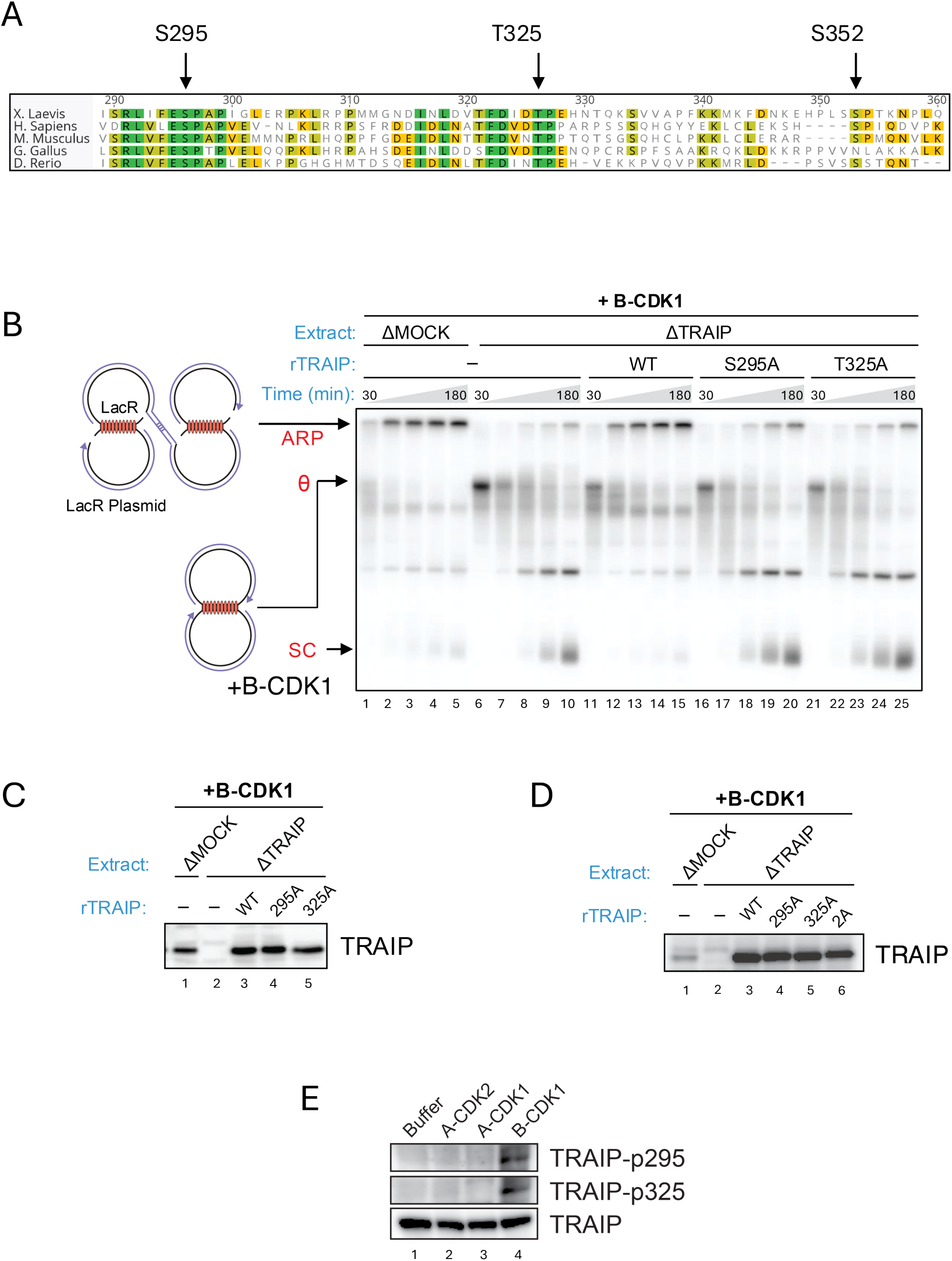
TRAIP phosphorylation and function in Xenopus egg extracts. (A) Multiple sequence alignment showing conserved CDK consensus sites in TRAIP (Xenopus residue numbers shown above alignment). (B) Formation of aberrant replication products (ARP) during replication of a LacR plasmid in Xenopus egg extracts. The plasmid was replicated in mock-depleted or TRAIP-depleted extracts supplemented with B-CDK1 and the indicated recombinant TRAIP proteins. Samples from the indicated time points were analyzed by native agarose gel electrophoresis and autoradiography. ARP, θ structures (stalled replication forks), and SCs (supercoiled replication products) are indicated. (C) Immunoblot showing the relative amounts of recombinant TRAIP proteins added to B-CDK1-containing, TRAIP-depleted extracts used for the experiment shown in Fig. 1E. (D) Immunoblot analysis of recombinant TRAIP proteins added to TRAIP-depleted extracts used for the experiment shown in Fig. 1F-G. (E) Phosphorylation of TRAIP by different cyclin-CDK complexes. NPE was supplemented with the indicated CDK complexes (500 nM Cyclin A2-CDK2, 500 nM Cyclin A2-CDK1, or 300 nM Cyclin B-CDK1) and analyzed by immunoblotting using phospho-specific TRAIP antibodies.

**Fig. S3.**
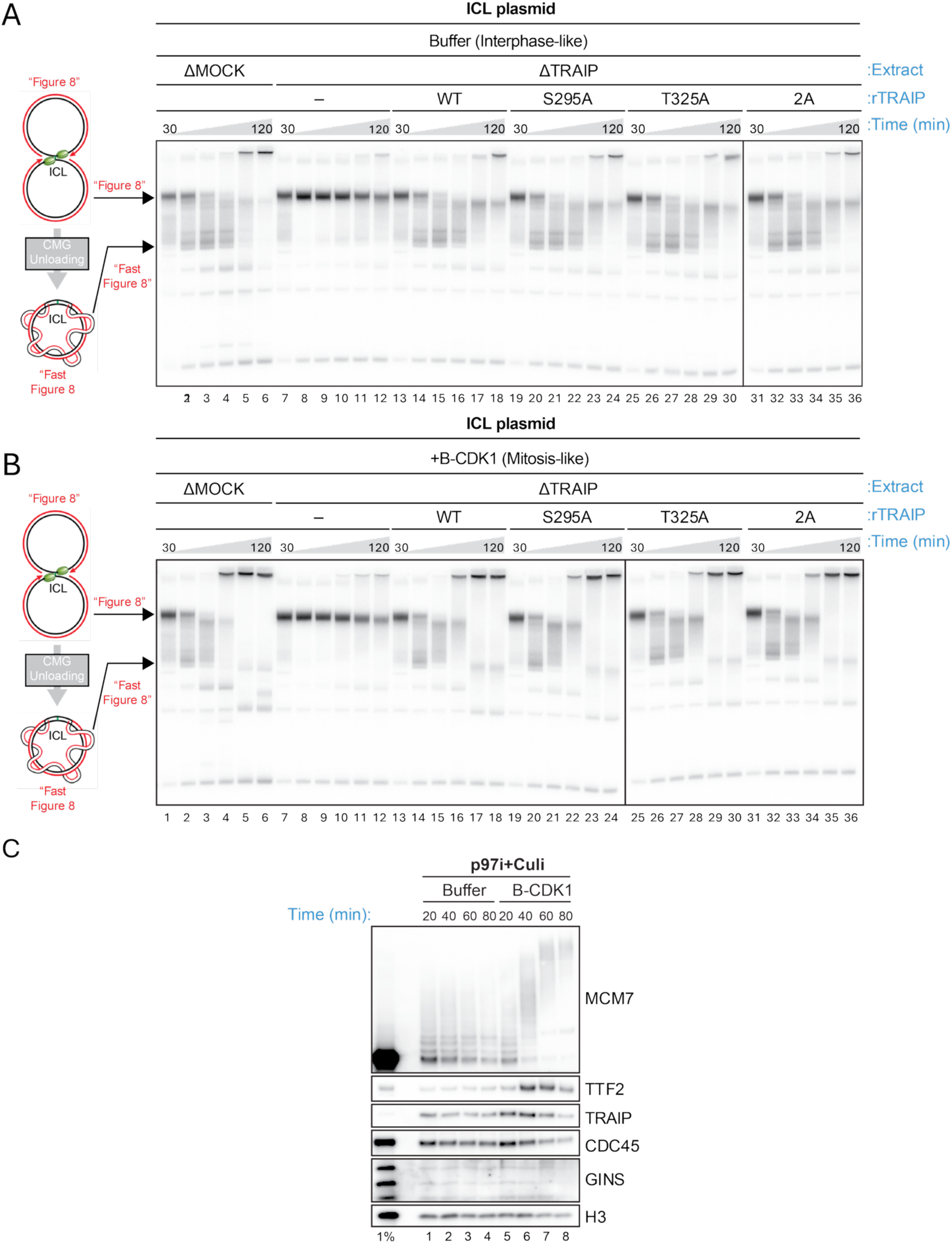
TRAIP phospho-site mutants support unloading of CMGs converged at ICLs in interphase and mitotic extracts. (A) Replication of an interstrand crosslink (ICL)-containing plasmid in interphase Xenopus egg extracts. The plasmid was replicated in mock-depleted or TRAIP-depleted extracts supplemented with the indicated recombinant TRAIP proteins. Samples from the indicated times were collected and analyzed by native agarose gel electrophoresis and autoradiography. Figure 8: converged CMGs at an ICL; Fast Figure 8: similar species as Figure 8, but after CMG unloading, which induces a faster-migrating, catenated species (58). (B) The experiment from (A) was performed in B-CDK1 containing extracts. (C) ICL-containing plasmid was pulled down from the indicated extracts to assess MCM7 ubiquitylation and the retention of the indicated proteins. Replisome unloading was blocked by p97i and Culi.

**Fig. S4.**
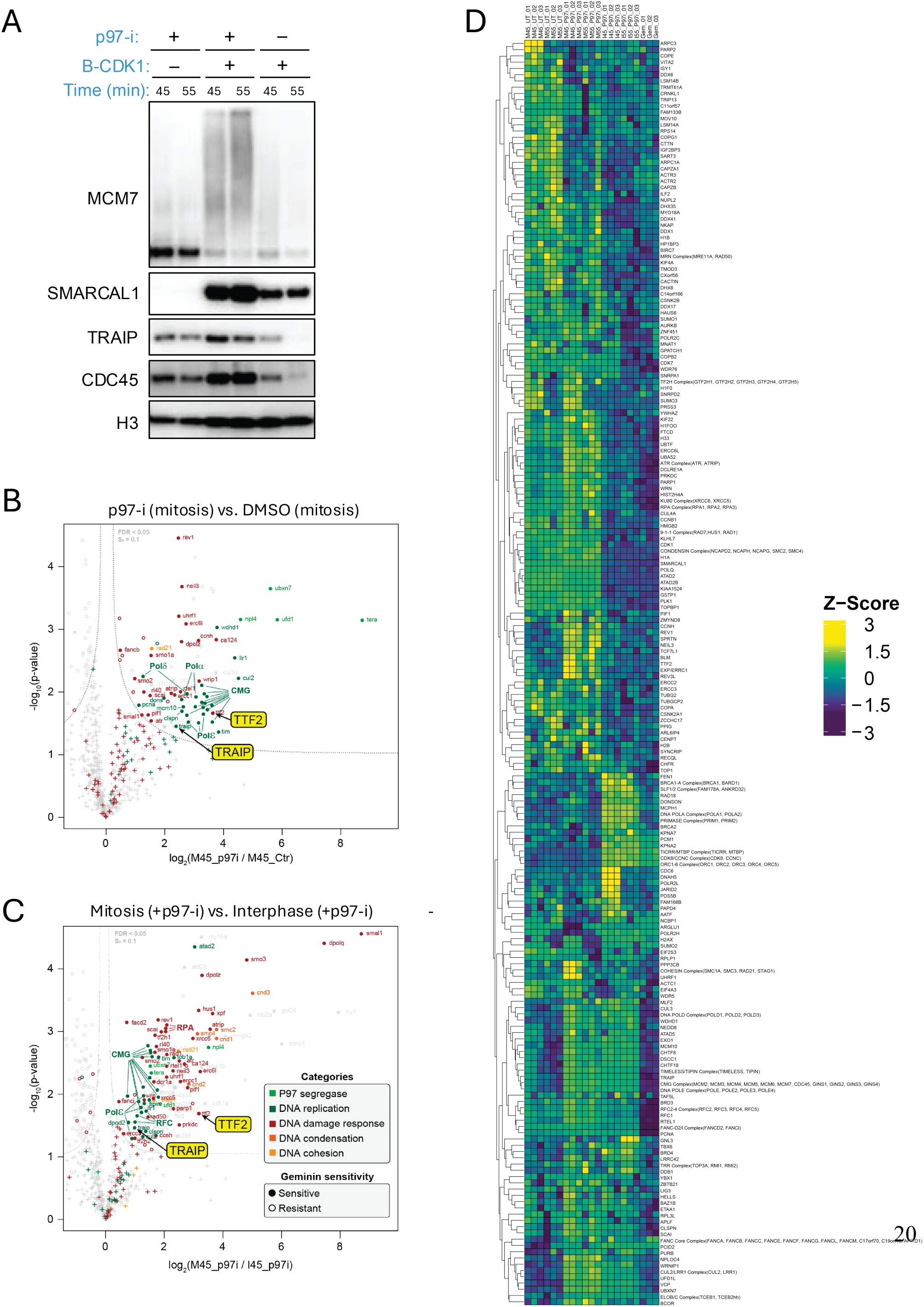
Mass spectrometry analysis of proteins associated with mitotic replisomes. **(A)** LacR plasmid was replicated in the indicated egg extracts. Samples were recovered 45 and 55 minutes after NPE was added to initiate replication. Retention of the indicated proteins was assessed by western blotting. **(B)** Sample from (A) plus two additional replicates were analyzed by mass spectrometry. The heatmap represents mass spectrometry results showing genome maintenance protein abundance. Protein levels are shown as the mean of the Z scored log2 abundance in the different conditions. M, mitotic extract; I, interphase extract; UT, untreated; p97i, p97 inhibitor; Gem, geminin-treated, a control without replication. In addition to individual proteins, some protein complexes are shown as the integrated abundance over their subunits. Full results are reported in Table S1. **(C)** Volcano plot of protein abundance from (B) in the presence and absence of p97-i (both containing B-CDK1; 45 min timepoint). The plot displays the mean difference of the protein intensity (-log_2_) versus the p value (-log_10_) calculated by a modified, two-sided t test. **(D)** Same as (C), except the volcano plot compares recruitment in the presence and absence of B-CDK1 (both conditions contain p97-i; 45 min time point).

**Fig. S5.**
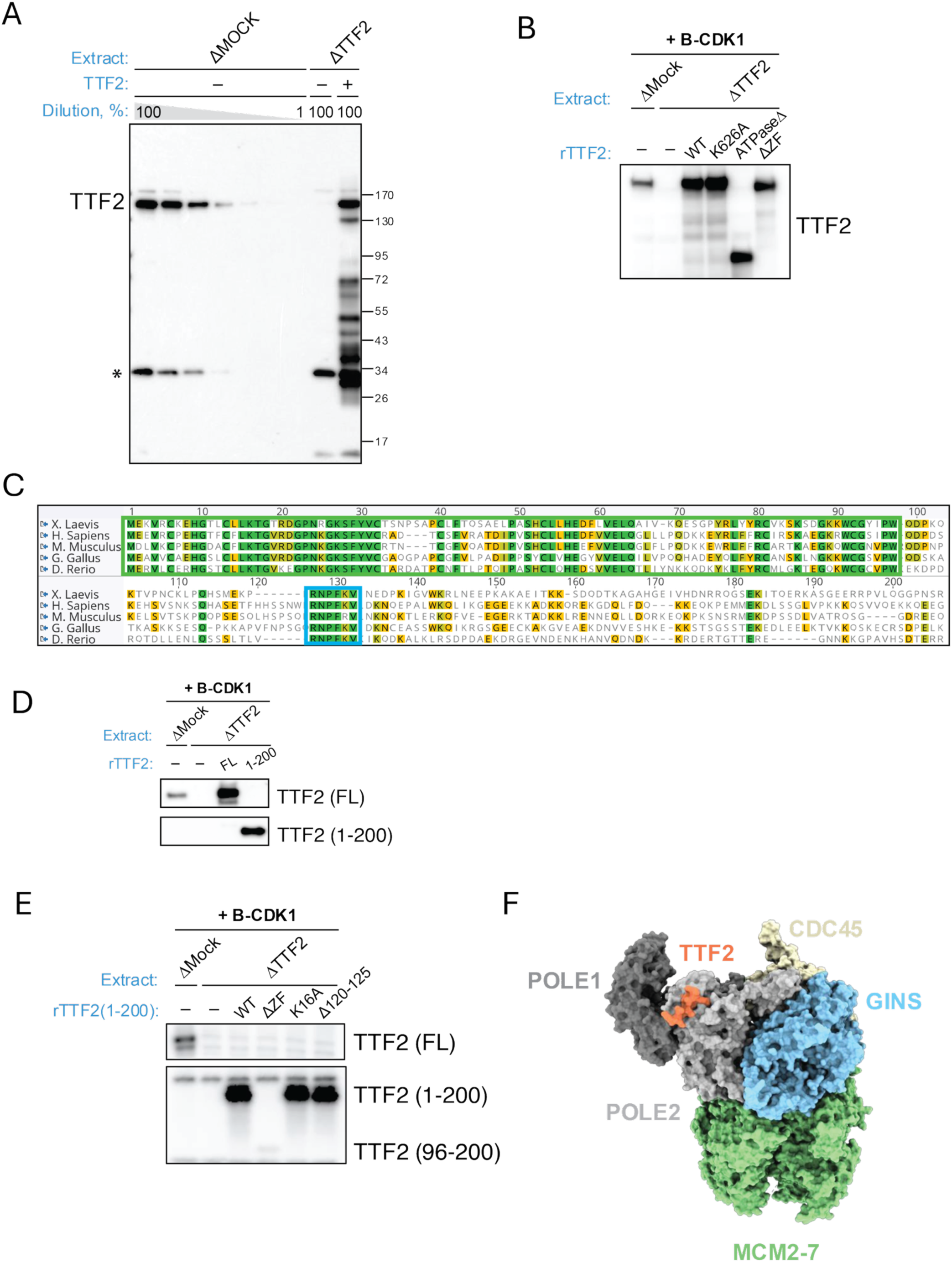
TTF2 antibody characterization and expression of recombinant TTF2 and fusion constructs. **(A)** High speed supernatant (HSS) of egg lysate was mock-depleted, depleted of TTF2, or depleted of TTF2 and supplemented with recombinant TTF2 expressed by in vitro transcription-translation (TTE). Extracts (including a dilution series of mock-depleted extract) were blotted with an antibody raised against *Xenopus* TTF2. *, presumed cross-reacting protein. (**B)** Extracts from Fig. 2D-F. NPE was mock-depleted or depleted of TTF2, and transcription-translation extract (TTE) programmed with empty vector or the indicated TTF2 constructs was added. The NPE-TTE mixture was blotted for TTF2. **(C)** Multiple sequence alignment showing the N-Terminus (residues 1-200) of TTF2. The conserved Zinc fingers domain is shown in green box and conserved POLE2 binding motif is shown in blue box. **(D)** Similar to (B) but corresponding to Fig. 2G. **(E)** Similar to (B) but corresponding to Fig. 3B-D. TTF2^1-200/ΔΖF^ is barely detectable, which could be due to poor expression or poor reactivity with the TTF2 antibody, which is raised against a larger fragment of the TTF2 N-terminus. **(F)** TTF2-binding to POLE2 should not sterically clash with other replisome proteins. The predicted POLE2-TTF2 complex from Fig. 3E was aligned on POLE2 in the replisome structure (PDB: 7PLO).

**Fig. S6.**
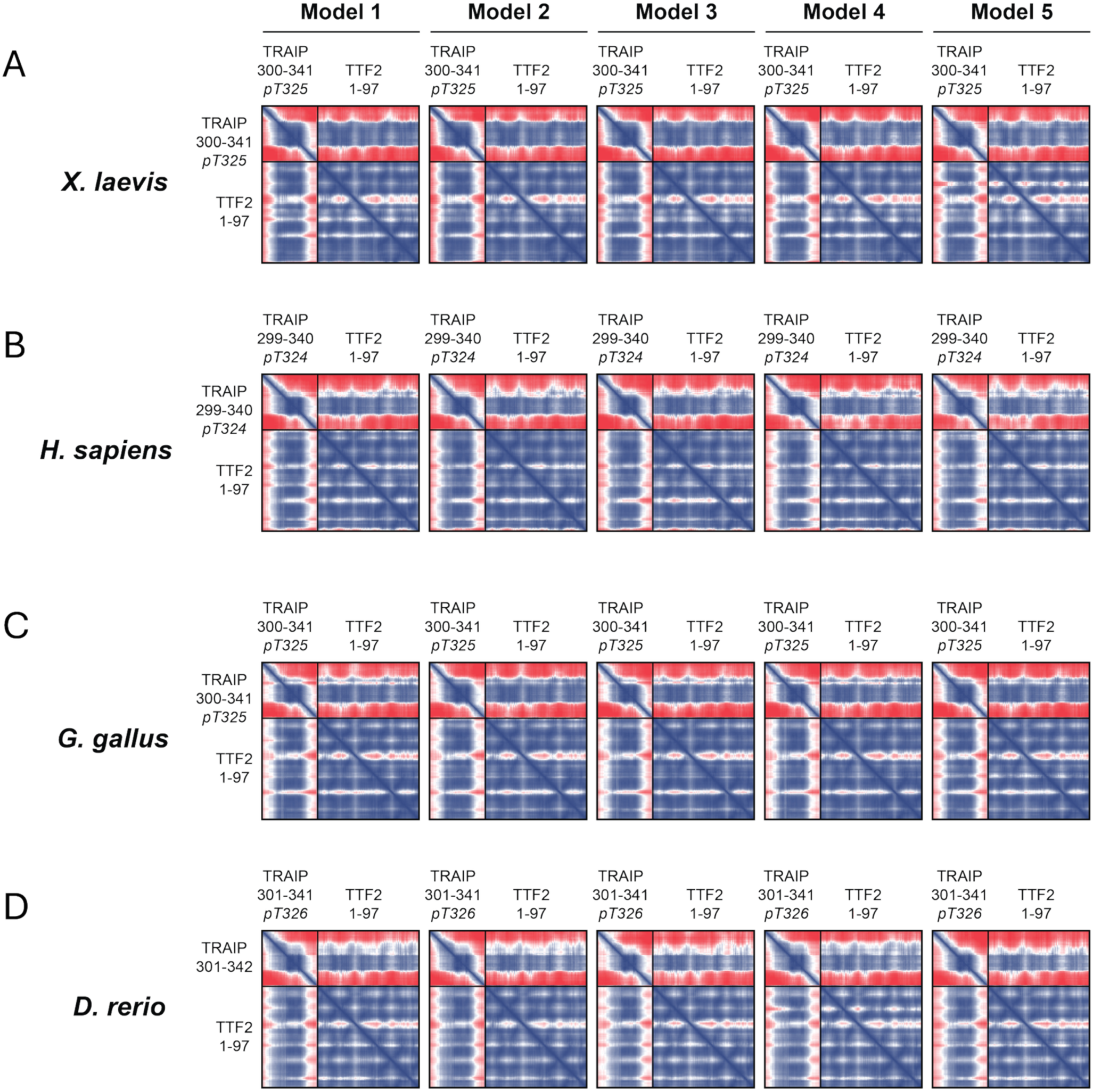
AlphaFold3 predicts a conserved phospho-dependent TRAIP–TTF2 interaction. Predicted aligned error (PAE) plots from AlphaFold3 modeling of the TRAIP C-terminal region containing the phosphorylated CDK site and the N-terminal region of TTF2 (residues 1–97). Lower PAE values (blue) indicate higher confidence in the relative positioning of residues and therefore greater confidence in the predicted interaction interface. **(A)** Homo sapiens TRAIP (299–340, pT324) with TTF2(1–97). **(B)** Xenopus laevis TRAIP (300–341, pT325) with TTF2(1–97)**. (C)** Gallus gallus TRAIP (300–341, pT325) with TTF2(1–97)**. (D)** Danio rerio TRAIP (301–342, pT326) with TTF2(1–97). Across species, low PAE is observed at the interface between phosphorylated TRAIP and the N-terminal zinc finger region of TTF2, supporting a conserved phospho-dependent TRAIP–TTF2 interaction.

**Fig. S7.**
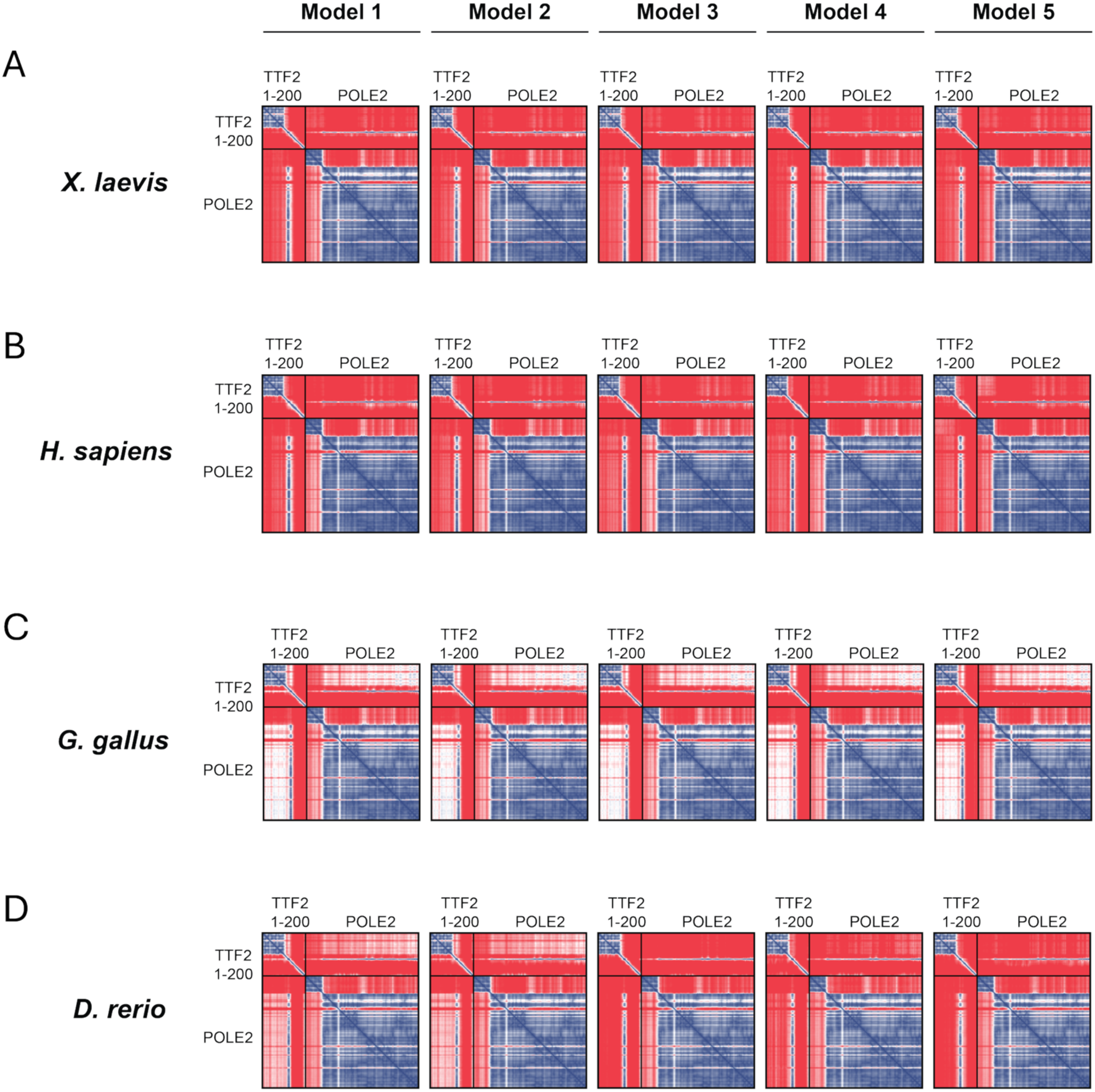
AlphaFold3 predicts a conserved TTF2-POLE2 interaction. Predicted aligned error (PAE) plots for AlphaFold3 models of TTF21-200 in complex with the POLE2 subunit of DNA polymerase ε from the same organism. Lower PAE values (blue) indicate higher confidence in the relative positioning of residues and therefore greater confidence in the predicted interaction interface. Across species, the conserved motif adjacent to the TTF2 zinc fingers displays a consistently low PAE with POLE2, supporting a conserved interaction between TTF2 and the Pol ε complex. **(A)** Homo sapiens TTF21-200 with POLE2. **(B)** Xenopus laevis TTF21-200 with POLE2. **(C)** Gallus gallus TTF21-200 with POLE2. **(D)** Danio rerio TTF21-200 with POLE2.

**Fig. S8.**
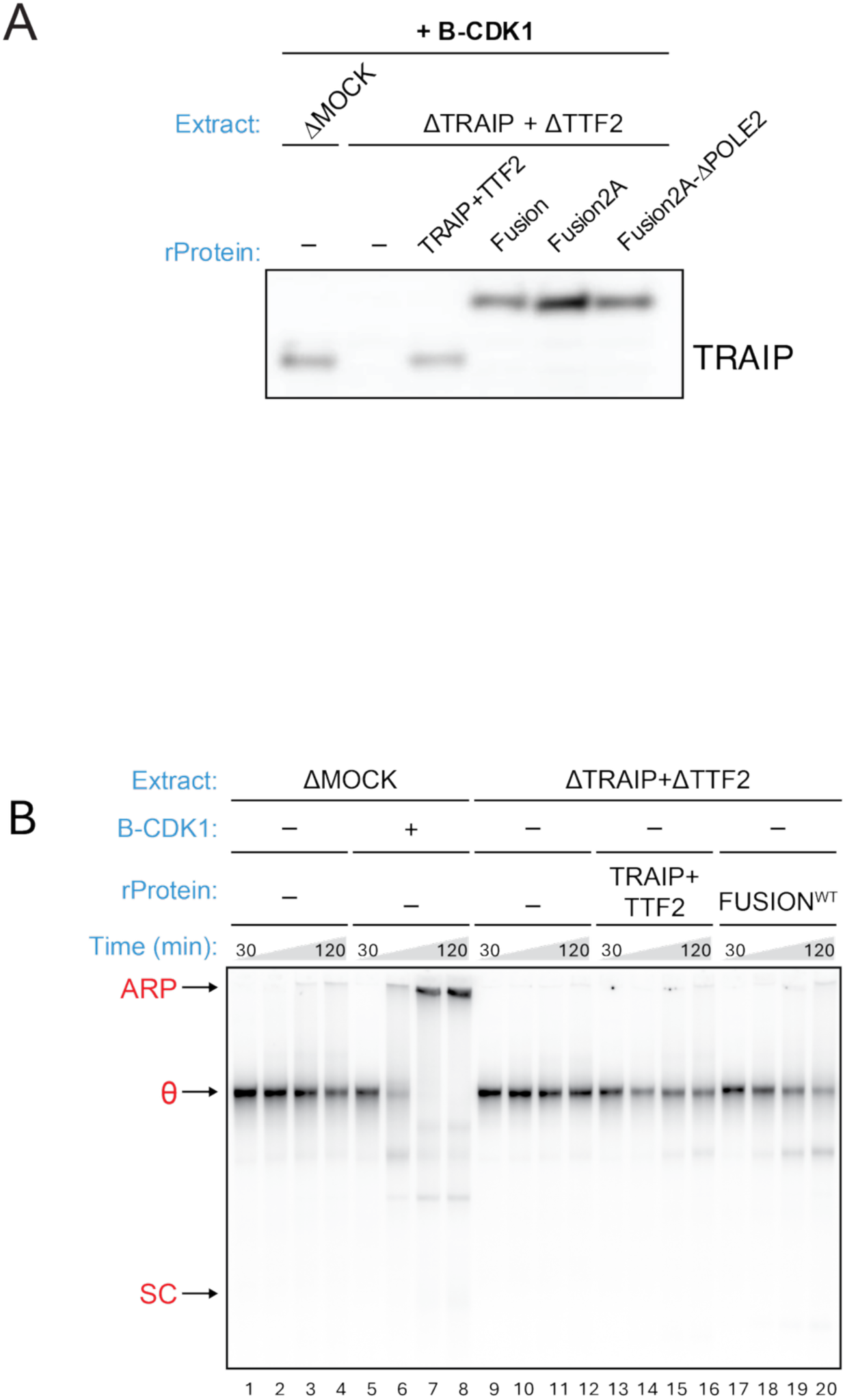
The TRAIP–TTF2 fusion does not trigger ARP formation in interphase extract. **(A)** Western blot showing the indicated TRAIP proteins added to the indicated extracts. **(B)** LacR plasmid replication assay performed in Xenopus egg extract under the indicated conditions. Extracts were either mock-depleted (ΔMOCK) or depleted of both TRAIP and TTF2 (ΔTRAIP+ΔTTF2). Where indicated, recombinant TRAIP and TTF2 or the TRAIP–TTF2 fusion (FUSIONWT) proteins were added. B-CDK1 was added as indicated. Samples were collected at 30 and 120 minutes after initiation of replication and separated on a native agarose gel. Positions of aberrant replication products (ARP), theta structures (θ), and supercoiled plasmid (SC) are indicated. In mock-depleted extract supplemented with B-CDK1, ARP formation was observed (lanes 6-8). In contrast, in TRAIP and TTF2 depleted extract lacking B-CDK1 (interphase condition), addition of TRAIP and TTF2 or FUSIONWT did not promote ARP formation or loss of θ structure plasmids with stalled replication forks.

**Fig. S9.**
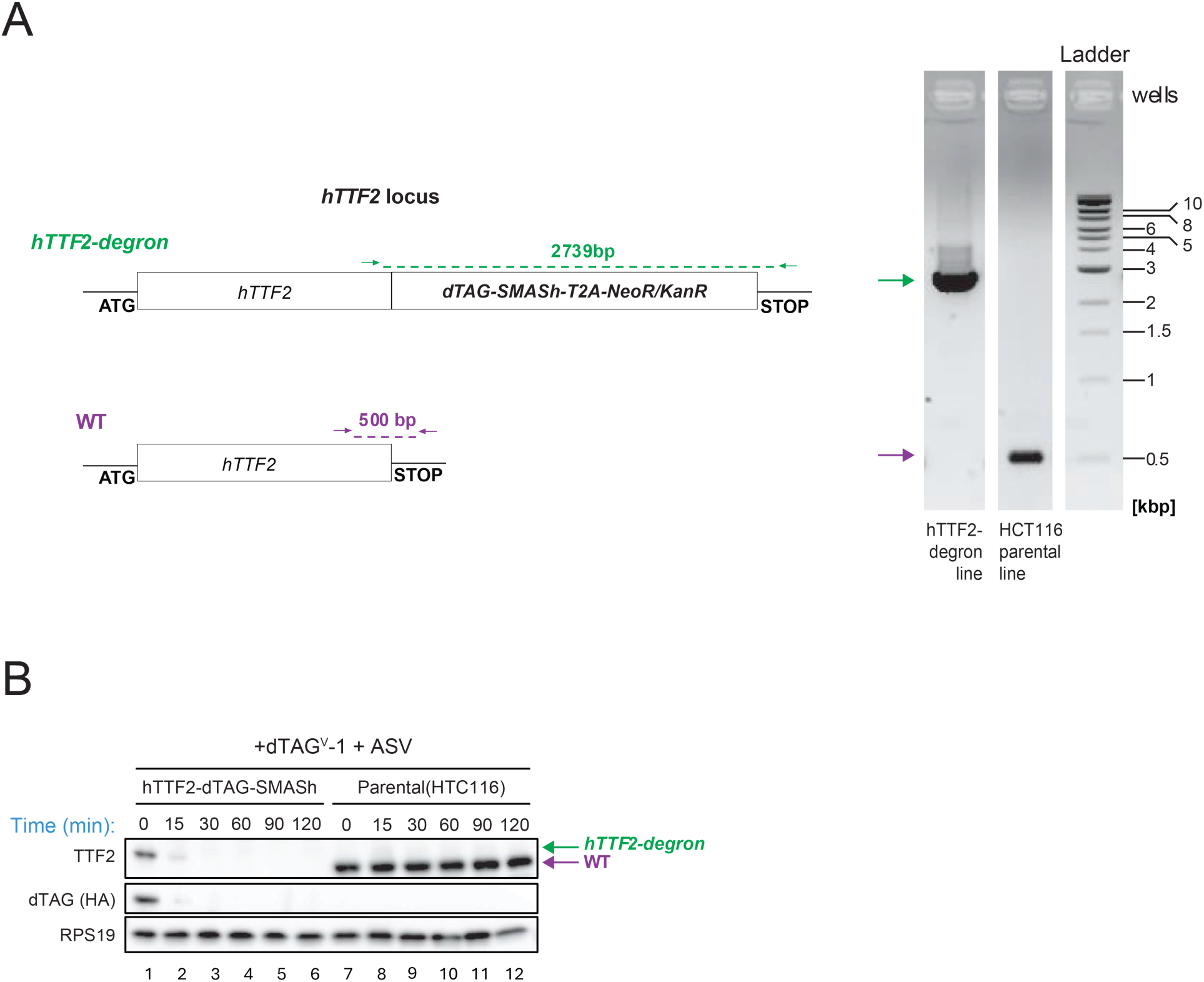
Generation and validation of the human TTF2 degron cell line. **(A)** The STOP codon of the human TTF2 gene was replaced with a sequence encoding two tandem degron tags (dTAG-SMASh), using CRISPR-Cas9 genome editing (left). Clones with the desired modification were identified by PCR genotyping (right). **(B)** Conditional degradation was achieved by concomitant addition of dTAG^V^-1 (to initiate degradation of the dTAG tag) and ASV (to initiate degradation of the SMASh tag). The degron-tagged TTF2 protein is degraded within 30 minutes of ligand addition.

**Fig. S10.**
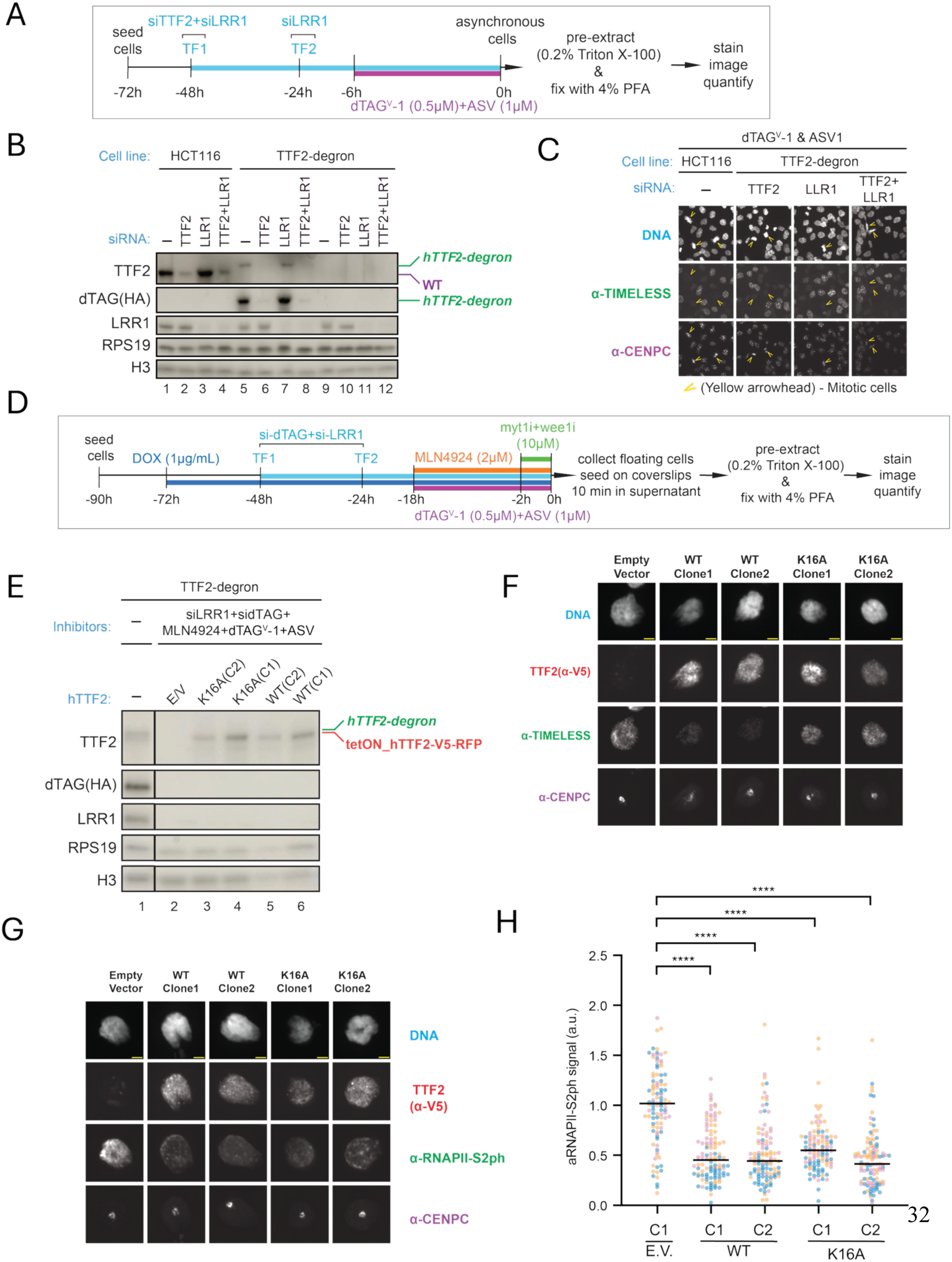
Assay for mitotic CMG unloading in human cells. **(A)** Schematic of experimental workflow used to assess TTF2 contribution to TIMELESS removal from chromatin. Cells were seeded in duplicates on 12-well plates (the first plate without coverslips, the second with) and treated identically. At 0-hour time point, cells from the first plate were trypsinized to prepare total protein extracts for immunoblotting, shown in (B), while cells on the second plate (with coverslips) were pre-extracted, fixed, and stained, shown in (C). **(B)** Western Blots to test TTF2 and LRR1 protein depletion under conditions as in (A). **(C)** Representative images of fixed and stained cells under conditions as in (A). **(D)** Schematic of the experimental workflow used to assay TIMELESS retention on mitotic chromatin under conditions of stringent CRL2^Lrr1^ and TTF2 inhibition. To downregulate CRL2^Lrr1^, an siLRR1 based double transfection (TF1 and TF2) was combined with Cullin-RING E3 ligase (CRL) inactivating neddylation inhibitor MLN4924. To downregulate TTF2 under these conditions of decreased CRL activity (itself required for dTAG-based degradation), we combined dTAG^V^-1 and ASV with a double siRNA transfection targeting the dTAG part of endogenous TTF2-degron (which does not target the add-back TTF2 constructs). Cells were treated with doxycycline (DOX) to induce the expression of the add-back constructs, and with a combination of WEE1 and MYT1 kinase inhibitors to induce premature mitotic entry. Mitotic cells were then collected, attached to coverslips, pre-extracted, fixed, stained, imaged, and the signal intensities were quantified, and shown in (F, G, H, Fig. 5D). A sample of collected mitotic cells was used to prepare total protein extracts for immunoblotting, shown in (E). **(E)** Western Blotting to test for TTF2 and LRR1 protein depletion under stringent conditions as in (D). **(F)** Representative images of TIMELESS retention on mitotic chromatin under conditions as in (D), quantification is shown in (Fig. 5D). Mitotic cells expressing empty vector, TTF2^WT^, or TTF2^K16A^ clones were stained for DNA, TTF2 (α-V5), TIMELESS, and CENPC. **(G)** Representative images of RNAPII-S2ph retention on mitotic chromatin under the same conditions as in (D). **(H)** Quantification of RNAPII-S2ph on mitotic chromatin as in (G). Three experimental repeats are shown, 40 cells per sample in each with medians as horizontal lines. Mann-Whitney test was used to assess the significance of differences (****, p < 0.0001).

**Fig. S11.**
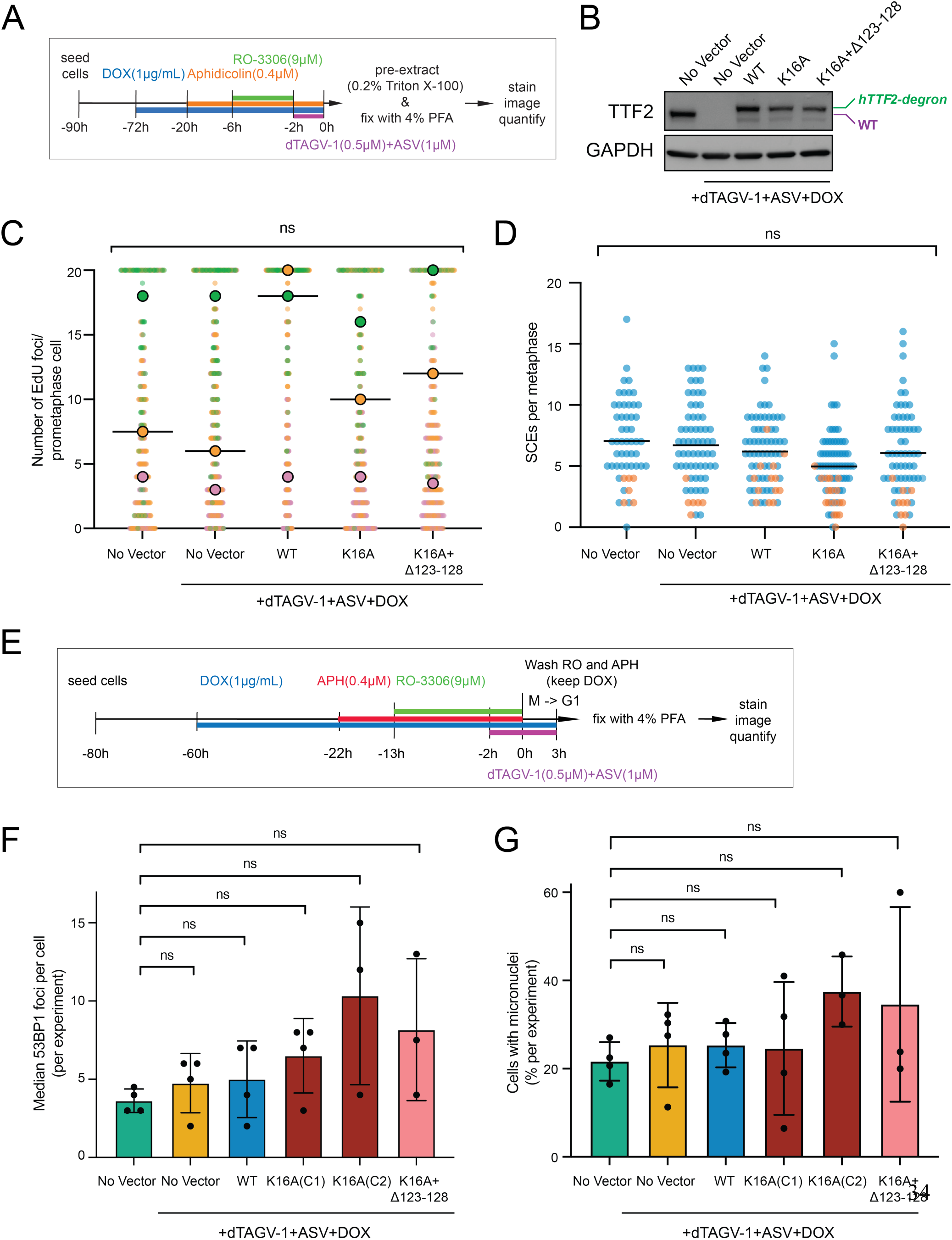
Disruption of the TRAIP–TTF2 interaction does not strongly affect MiDAS, sister chromatid exchange, or other markers of genome instability. **(A)** Schematic of the experimental setup used in MiDAS and SCE assays, except BrdU added 48h before harvest for the latter. Briefly, cells were induced with doxycycline (dox) and aphidicolin. The cells were arrested in G2 with CDK1 inhibitor (RO-3306) and released into mitosis with EdU. Mitotic cells were collected, pre-extracted, fixed, stained, imaged, and quantitated. **(B)** Representative western blots showing the degradation of endogenous TTF2 and expression of complemented constructs and GAPDH as loading control. **(C)** Distribution of EdU foci per individual cell under the indicated complementation conditions (+dTAG^V^-1 + ASV + DOX). Each dot represents the number of EdU foci per prometaphase figure; horizontal bars indicate the median. Degradation of TRAIP similarly showed no effect on EdU foci distribution (data not shown) **(D)** Distribution of SCEs per individual metaphase cell under the indicated conditions (+dTAG^V^-1 + ASV + DOX). Each dot represents one metaphase; horizontal bars indicate the median. Statistics in panels C and D were performed using two-tailed, non-parametric Mann-Whitney tests. For panel C, the medians of each condition were compared. **(E)** Schematic of the experimental setup used in 53BP1 foci and micronuclei formation. **(F)** Quantification of 53BP1 foci per nucleus (median per experiment) under the indicated treatment conditions. Cells were treated with dTAG^V^-1 and ASV to degrade endogenous TTF2 and complemented with the indicated constructs in the presence of aphidicolin (APH). Bars represent mean ± s.d.; dots represent independent experiments. **(G)** Percentage of cells with micronuclei under the indicated conditions (± dTAG^V^-1 + ASV; + APH). Bars represent mean ± s.d.; dots represent independent experiments. Wilcoxon matched-pairs signed rank test was used to compare distribution of medians between TTF2^WT^ (no degradation, first column) and the rest of samples, all differences were not significant (F, G).

**Fig. S12.**
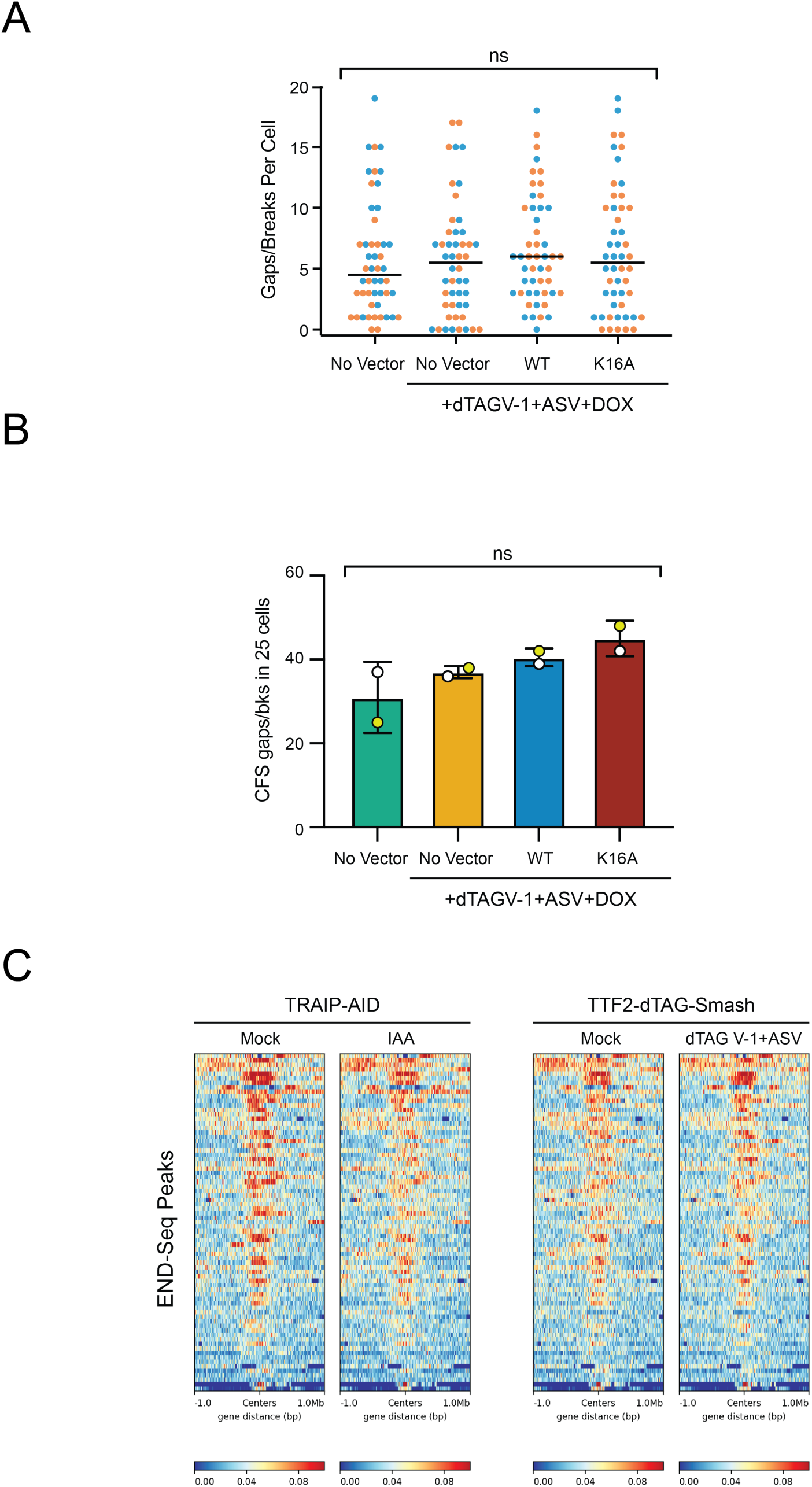
Fragile site expression and genome-wide DNA break mapping upon disruption of the TRAIP-TTF2 complex. **(A)** Quantification of total chromosomal gaps/breaks per cell in HCT116 TTF2-degron cells complemented with the indicated constructs (No vector, WT, K16A) under degron conditions (+dTAG^V^-1 + ASV + DOX) under experimental setup as in (fig. S11A). Each dot represents one metaphase; horizontal bars indicate the mean. **(B)** Frequency of breaks at fragile sites FRA3B and FRA16D in HCT116 TTF2-degron cells complemented with the indicated constructs (No vector, WT, K16A) under degron conditions (+dTAG^V^-1 + ASV + DOX). Error bars indicate 95% confidence level. For panels A and B, significance was determined using ordinary one-way ANOVA. **(C)** Heatmaps of END-seq signal centered on peak regions under the indicated conditions. Left, TRAIP-AID cells treated with mock or IAA. Right, TTF2-dTAG-SMASh cells treated with mock or dTAG^V^-1 + ASV. Signal intensity is plotted across ±1 Mb from peak centers. Color scale indicates normalized read density.

**Fig. S13.**
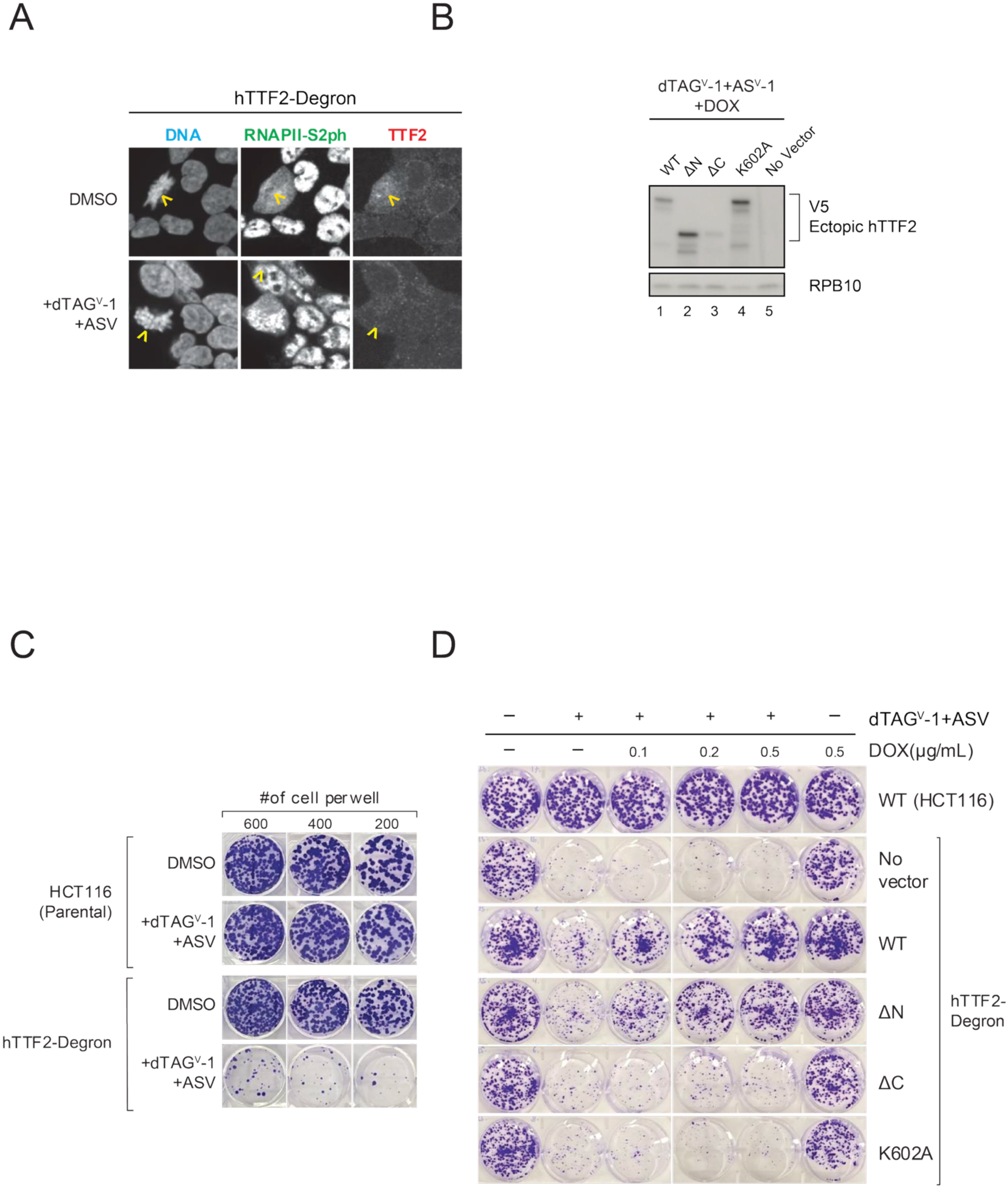
RNAPII eviction requires the TTF2 ATPase domain and correlates with cell proliferation. **(A)** TTF2 degradation leads to retention of phosphorylated RNAPII on mitotic chromosomes. hTTF2-degron cells were treated with dTAG^V^-1 and ASV as indicated and stained for DNA, RNAPII-S2ph, and TTF2. Yellow arrowheads indicate mitotic cells. **(B)** Immunoblot detection of endogenous hTTF2-degron depletion and induction of ectopic tetON complementation constructs. Cells expressing the indicated hTTF2-V5-RFP constructs were treated with dTAG^V^-1, ASV, and DOX and blotted with antibodies against hTTF2, V5, and RPS19. Orange arrowheads point to the location of hTTF2-V5-RFP species on the blot. **(C)** Clonogenic proliferation assay comparison of parental HCT116 cells and the hTTF2-degron line in the presence or absence of dTAG^V^-1 and ASV. Representative crystal-violet-stained wells are shown. **(D)** Clonogenic proliferation assay for complementation of the growth defect after endogenous TTF2 degradation with the indicated tetON hTTF2 constructs. Cells were treated with dTAG^V^-1 and ASV and induced with the indicated DOX concentrations. Representative plates are shown for no vector, TTF2 WT, ΔN, ΔC, and K602A complementing constructs.

